# ARZIMM: A Novel Analytic Platform for the Inference of Microbial Interactions and Community Stability from Longitudinal Microbiome Study

**DOI:** 10.1101/2022.01.26.477892

**Authors:** Linchen He, Chan Wang, Jiyuan Hu, Zhan Gao, Emilia Falcone, Steven Holland, Martin J. Blaser, Huilin Li

## Abstract

Dynamic changes of microbiome communities may play important roles in human health and diseases. The recent rise in longitudinal microbiome studies calls for statistical methods that can model the temporal dynamic patterns and simultaneously quantify the microbial interactions and community stability. Here, we propose a novel autoregressive zero-inflated mixed-effects model (ARZIMM) to capture the sparse microbial interactions and estimate the community stability. ARZIMM employs a zero-inflated Poisson autoregressive model to model the excessive zero abundances and the non-zero abundances separately, a random effect to investigate the underlining dynamic pattern shared within the group, and a Lasso-type penalty to capture and estimate the sparse microbial interactions. Based on the estimated microbial interaction matrix, we further derive the estimate of community stability, and identify the core dynamic patterns through network inference. Through extensive simulation studies and real data analyses we evaluated ARZIMM in comparison with the other methods.

## 1 Introduction

The human microbiota, a diverse array of microbial organisms living in and on human bodies, form a dynamic ecosystem that plays a critical role in human health. While temporally stable microbial communities are observed among healthy adults [1], the fluctuation of microbiome has been linked to increasing frailty [2] and declining immune function of hosts [3], and diseases such as inflammatory bowel disease [4, 5], colorectal cancer [6, 7], and irritable bowel syndrome [8, 9]. When a microbial community changes in response to an external perturbation, it undergoes a dynamic process and tends to evolve toward another stable state (Figure 1). This dynamic process is stochastic and varies according to the type and strength of perturbation, the community stability prior to the perturbation, and other subject-level relevant features. The recent rise in longitudinal studies, in which microbial samples are collected repeatedly over time, offers unique insights into the responses of such communities to perturbations and the associated dynamic patterns. For example, in our ongoing microbiome study evaluating the effects of antibiotic exposure as a short-term perturbation on microbial, immune, and metabolic physiology (MIME study), we are interested in determining how differently the microbial community responds to the antibiotic treatment.

**Figure 1:**
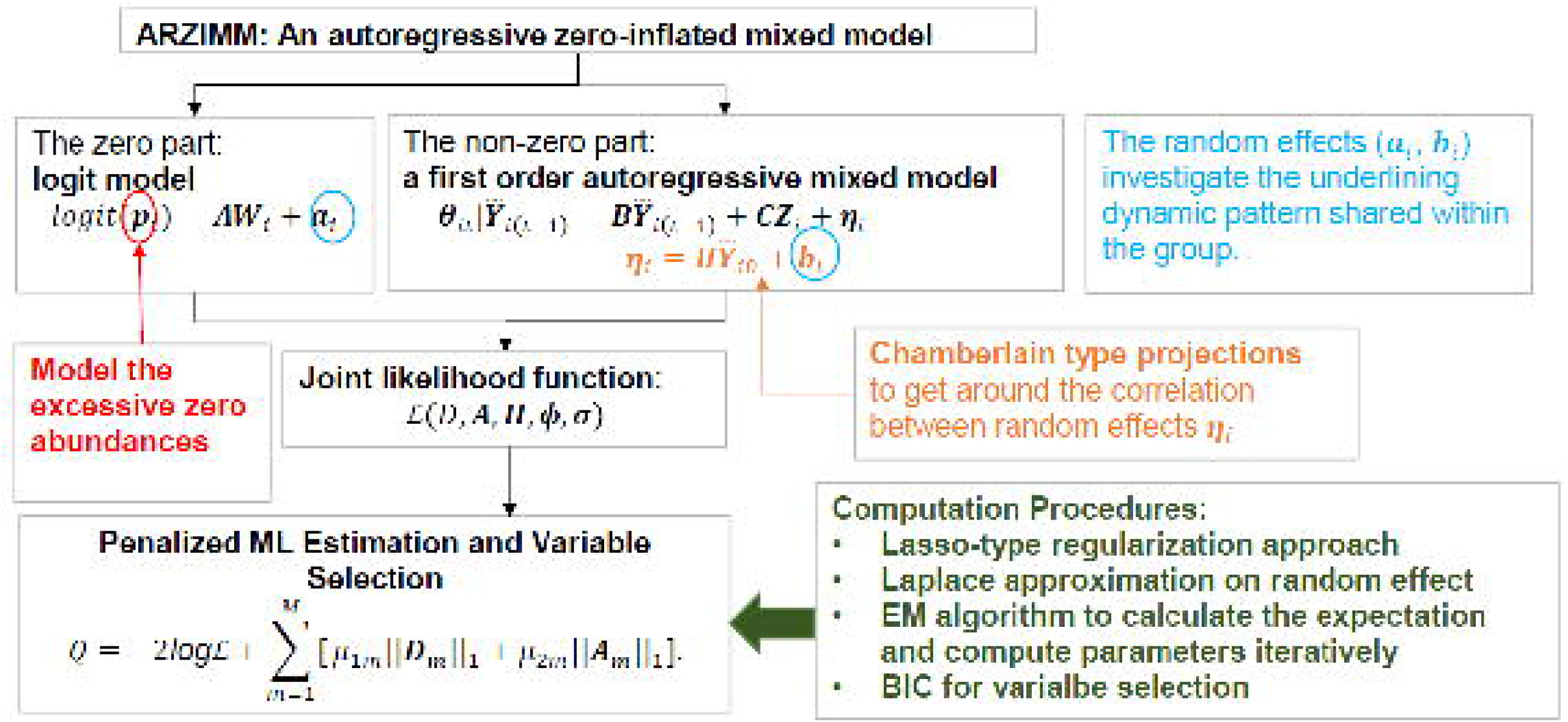
Schematic of the evolution of microbial community states in response to external perturbation. External perturbation (blue arrows) can affect microbial community composition (shown in a pie chart), defined as a community state. For each state, the ball-in-basin diagram portrays stability measured by the variance in the stationary distribution of the location of the ball. White arrows indicate the reaction of microbial community to the perturbation.

Human microbiota studies have been accelerated by the advent of next-generation sequencing technologies which enabled the quantification of the composition of microbiomes, often by two common sequencing approaches—16S rRNA marker gene sequencing and shotgun metagenomics sequencing [10]. There are pros and cons to each of those techniques, which are discussed in recent reviews [11, 12]. But for both methods, because of the varying sequencing read counts obtained across samples, it is necessary to employ various normalization tools to convert raw counts data into relative abundances [13]. However, the dependency of the compositional components greatly hampers the interpretation of microbiota changes in longitudinal studies. There is reason to believe that the absolute abundances of bacteria are biologically meaningful measures, especially in the study of microbial interactions. Thus, in our MIME study, we use an independent quantitative polymerase chain reaction (qPCR) technology [14-16] to quantify total bacterial load per unit sample, and then use these data to estimate absolute bacterial abundance by combining them with the relative abundance values obtained from 16S rRNA or shotgun sequencing methods. This MIME study motivated us to develop analytical methods to investigate microbial interaction and community stability after a strong external perturbation, and identify core active microbial taxa by modeling the absolute abundances of bacteria.

Although many well-developed statistical tools are widely used for assessing the diversity of microbial communities and its composition, there are only a few methods available for inferring the ecological networks of microbial communities. Here we briefly review the well-developed statistical methods for studying the dynamic microbial systems and their limitations.

A Bayesian network contains a set of multivariate joint distributions that exhibit certain conditional independences and a directed and acyclic graph (DAG) that encodes conditional independences among random variables. If the dependence relationships repeat and the signals at a certain time point only depend on the signals from previous time points, then the whole network can be formulated as a dynamic Bayesian network (DBN)[17] representation. McGeachie et al.[18] constructed a simplified two-stage DBN (TS-DBN) which uses a Markov assumption that the observed values at time *t* + 1 are independent of those at earlier time points (*t* − 1 and earlier) given the variable values at time *t*. Lugo-Martinez et al. presented a computational pipeline which first aligns the data collected from all individuals, and then learns a dynamic Bayesian network from the aligned profiles[19]. However, DBN has several limitations in analyzing the longitudinal microbial data. (1) It can only model the microbial community subject-by-subject. (2) DBN cannot handle the exceeding zero structure of microbial counts. Most methods remove the taxa whose relative abundances exhibit zero entry (i.e., not present in a measurable amount at one or more of the measured time points) before the downstream analysis. (3) The assumed distributions are unrealistic. E.g. all continue variables are assumed to be normally distributed. (4) The computational cost is relatively high, since parent nodes are added sequentially for each bacterial node. Additionally, the maximum number of possible parents is imposed, which is not realistic. (5) Due to sampling and sequencing limitations, the compositionality bias in microbiome data may also cause inaccurate estimation of parameters. The existing methods ignore this compositionality bias, making parameter estimates difficult to interpret. (6) Irregular sampling time may also result in inaccurate parameter estimation. Therefore, it is advised to cautiously interpret the findings from DBN[20, 21].

The classical Lotka-Volterra equations has been used to model simple system such as two species in a predator-prey relationship, where the interactions are strictly assumed to be competitive. The generalized Lotka-Volterra (gLV) equations extend the classical predator-prey (Lotka-Volterra) equations, where the interacting species might have a wide range of relationships including competition, cooperation, or neutralism. Assuming that the interaction (or the effect) of one species with another can be modeled by the corresponding coefficient in the equation, gLV equations provide a framework to analyze and simulate microbial populations. Mounier et al. used the gLV equations to model the interaction between bacteria and yeast in a cheese microbiome[22]. Other microbiome studies further extended and implemented the gLV equations[23-26].

Many software are available for applying gLV modeling on microbial time series data, such as LIMITS[27], MetaMis[28], and MDSINE[29]. LIMITS and MetaMis can be implemented to construct microbial interactions using the longitudinal microbiome data from one subject. MDSINE can jointly analyze multiple time series, but requires Matlab programming. Web-gLV (http://web.rniapps.net/webglv) can be used for modeling, visualization, and analysis of microbial populations, but can only handle limited number of samples. In summary, there are several limitations of gLV in analyzing the longitudinal microbial data. (1) gLV based models capture the interactions using a single averaged effect, thus they are not well-suited for noisy data. (2) Some methods estimate almost all possible edges without incorporating variable selection techniques. (3) gLV estimates the growth rate of each taxon marginally, therefore, ignores the intrinsic dynamic correlations of the repeated measurements. (4) gLV does not account for random processes which forms essential part of any biological system. (5) With the increased number of species and time span of prediction, the simulation output is prone to numerical errors. For example, Web-gLV can only simulate a maximum of 10 species at a time for at most 100 time points. (6) As DBN, gLV is not suitable for sparse, compositional, and irregular sampled microbiome data.

In Ives et al. [30], the stability of a microbial community is determined by three key interrelated components of microbial community structure: diversity, species composition, and interaction pattern among species. They viewed the dynamics of a microbial community as a stochastic process and proposed to use a first-order multivariate autoregressive process (MAR (1)) time-series model to disentangle the effects of these three components on community stability and to estimate the stability properties of a community by estimating the strengths of interactions between species. This method is widely used to estimate the stability of ecosystems (e.g., lake, ocean) based on culture-dependent microbial data[31, 32]. Usually a few (four or five) key microbes are detected with high frequency in each ecosystem in time-series measurements over a long period, and their abundances are rarely zero. In contrast, our MIME study will yield microbiome data from approximately nine time points over half a year from 80 subjects in three groups in the complete study—a relatively smaller number of repeated microbiome samples but from a relatively larger number of microbial communities (subjects) than what would be the case for an ecosystem study. Moreover, the 16S rRNA sequencing and qPCR methods used in this study provide absolute abundances for a staggering number of taxa, which include a large number of zero values. Because the MAR modeling methods require the normality assumption, they are not appropriate for analyzing data from sequence-based longitudinal microbiome studies. Therefore, we propose an autoregressive zero-inflated mixed effects model (ARZIMM) to address the special features of data instead. Its novelties are threefold. First, we propose to use a zero-inflated Poisson autoregressive model to model the excessive zero abundances and the non-zero abundances separately. Second, the random effects in the proposed model can investigate the underlining dynamic pattern shared within the group. Third, the employment of regularization techniques and network inference in our model enables the identification of the core dynamic patterns. The proposed ARZIMM estimates the strength of interactions between taxa, which is required to estimate the stability properties of a community, and identify key active taxa efficiently by using all of the longitudinal sequencing data. ARZIMM has been implemented in an open-source software package (https://github.com/Hlch1992/ARZIMM), and provides a useful tool for formulating, understanding, and implementing longitudinal microbiome data analysis.

In the following Material and Method section, we introduce the ARZIMM framework, discuss the quantification of microbial stability based on the estimated microbial interaction matrix, and investigate the conditions under which there exist a strict-sense stationary distribution. Then in the Result section, we evaluate ARZIMM using extensive simulation studies to show that it outperforms the conventional methods, and apply ARZIMM to the MIME study to illustrate network visualization and inference. In the end, we conclude with Discussion section.

## 2. Material and Method

### 2.1. ARIZMM Model

As illustrated in Figure 2, ARZIMM can be considered as a two-part model which comprises a logistic component and an autoregressive component. To address zero inflation, we consider the zero-inflated mixture model because it assumes both sampling zeros (due to the low sequencing depth) and structural zeros (being truly absent) exist in the data. Specifically, the logistic component models the structure zeros of taxa in the samples, and the autoregressive component models the non-structure-zero abundances of the taxa under the assumption that the changes in abundances from time *t* − 1 to time *t* depend only on the observed abundances at time *t* − 1 and other time-independent covariates and the observed abundances before time *t* − 1 have no direct effect. Since the goal of ARZIMM is to characterize microbial interactions and community stability during a short period after a strong external perturbation like the antibiotic usage in our MIME study, we assume there are no other time-dependent factors exist to affect the microbial stability.

**Figure 2:**
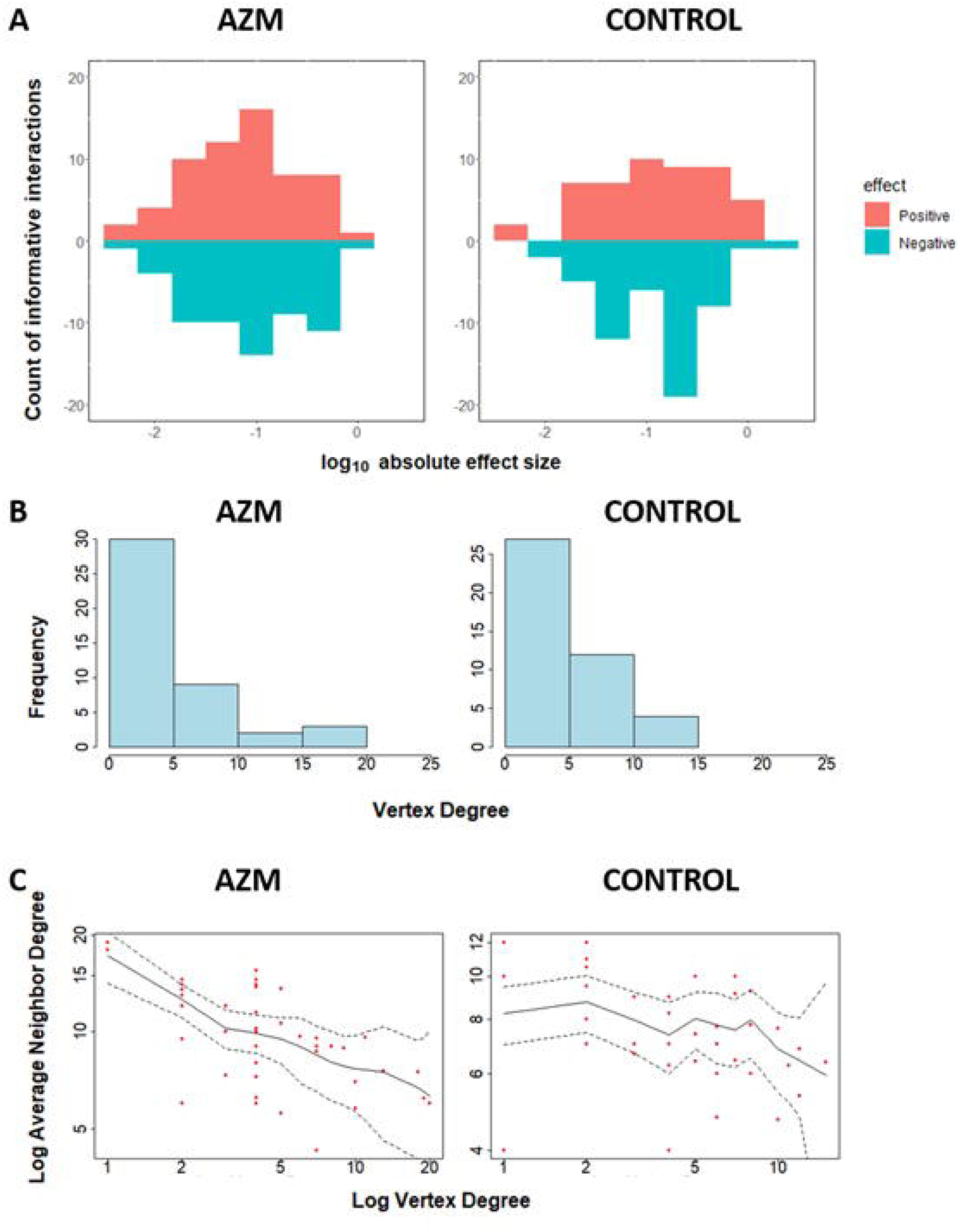
Graphical representation of ARZIMM model and analytic techniques.

### Notation and Model Specifications

Let *Y*_*imt*_ denote the observed absolute abundance of bacterial taxon *m* (*m* = 1, …, *M*) for subject *i* at time *t* (*i* = 1, 2, …, *n, t* = 1, …, *T*_*i*_), and we model *Y*_*imt*_ with a conditional mixture distribution as follow:

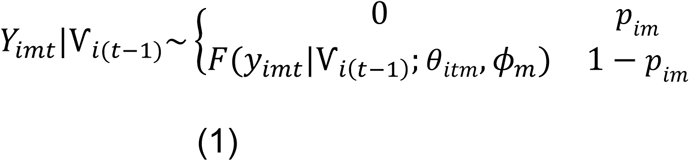

where *v*_*i*(*t*-1)_ represents all information that is known at time (*t* − 1) for individual *i*, including the observed absolute abundance *Y*_*im*(*t*-1)_ and later defined coviariates ***W***_*i*_ and ***Z***_*i*_. The parameter *p*_*im*_ represents the probability of the observation *Y*_*imt*_ being structural zero and is assumed time independent. Furthermore, *F* is assumed to be an exponential dispersion family distribution with the canonical parameter *θ*_*imt*_ and the dispersion parameter *ϕ*_*m*_. Both Poisson and negative binomial (NB) distributions can be used as to model absolute abundance. Below we illustrate the detailed modelling using Poisson model.

The mixture probability parameters ***p***_*i*_ = (*p*_*i*1_,…, *p*_*iM*_)′ are modeled by the logistic regression:

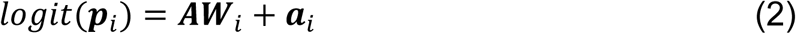

where ***W***_*i*_ = (1, *w*_*i*1_, …, *w*_*il*_)′ consists of intercept and *l* time independent covariates for individual *i*, the parameter ***A*** = (***A***_**1**_, …, ***A***_***M***_)′ is an *M* × (*l* + 1) matrix whose elements *A*_*mj*_ is the effect of covariate *j* on the zero proportion of taxon *m*. ***a***_*i*_ = (*a*_*i*1_, …, *a*_*iM*_)′ is an *M* × 1 vector of random intercepts to model the within-subject heterogeneity of being zero for individual *i* and has the joint multivariate normal distribution *𝒩*(**0, *∑***_a_).

The canonical parameters for Poisson distribution is *θ*_*imt*_ = log*E*(*Y*_*imt*_). We introduce the auto-regressive model by relating ***θ***_*it*_ = (*θ*_*i*1*t*_, …, *θ*_*iMt*_), to the *i*^*t*h^ individual’s observed log-transformed absolute abundance vector at time 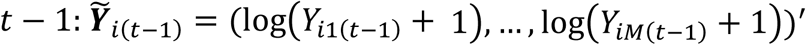. (where the pseudo count 1 is added to avoid the undefined logarithm when the absolute abudance is zero), and ***Z***_*i*_ = (1, *Z* _*i*1_, …, *Z* _*iq*_), the intercept and *q* time-independent covariates of individual *i* by

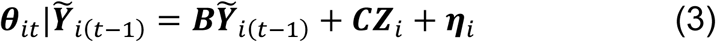

where ***B*** is an *M* × *M* matrix whose element *B*_*mj*_ gives the effect of the abundance of taxon *j* on the growth rate of taxon *m*, ***C*** is an *M* × (*q* + 1) matrix whose element *C*_*mj*_ gives the effect of covariate *j* on taxon *m*, and ***η***_*i*_ = (*η*_*i*1_, …, *η*_*iM*_)′ is time-independent random intercepts. Note that, as an autoregressive model, ***η***_*i*_ is correlated with the fixed effect 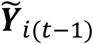 and this dependency can be tracked all the way back to the initial observation 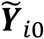. Because the standard random effects model has assumption that the random effects are independent to the other covariates in the model, in order to derive the random effect type maximum likelihood (ML) estimators, we use the Chamberlain type projections[33] to get around this correlation. Specifically, we project ***η***_*i*_ onto the time 0 observations 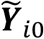 by:

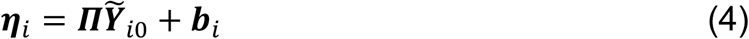

where ***Π*** is an *M* × *M* matrix with diag(***Π***) = (*π*_1_, …, *π*_*M*_)′ and off-diagonal components being zero. The components of ***Π*** represent how much variation in ***η***_*i*_ is due to the dependence on subject *i*’s initial value 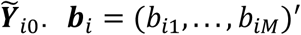 is an *M* × 1 vector, representing the independent subject-specific random effect and follows a joint multivariate normal distribution *𝒩*(**0, *Σ***_b_).

In the model, our primary interest is to estimate matrix ***B***, which measures the strengths of interactions between taxa. For a microbial community with a given number of species, its stability or dynamics status depends on the changes in the species’ population growth rates due to perturbation, which immediately cause the changes in the population growth rates of other species via species-species interactions[34]. Interaction between species can be viewed as a filter that amplifies the variability in species’ population growth rates caused by perturbation.

Note that we choose Poisson distribution because of its nice stationary distribution property in the autoregressive model which is crucial for our following stability investigation. To deal with the over-dispersion of microbiome data, we implemented the quasi-Poisson model [35] in the simulation and real data analysis.

### Penalized ML Estimation and Variable Selection

To define the joint likelihood of the longitudinal microbial absolute abundance data ***Y***_*it*_, we assume that the vector of time independent random effects 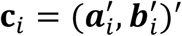 underlies both the zero and autoregressive generative processes and these random effects account for the within-subject group heterogeneity in the multivariate logistic component and the multivariate autoregressive component. Denote ***D*** = (***B, C***) = (***D***_**1**_, …, ***D***_***M***_),, ***ϕ*** = (*ϕ*_1_, …, *ϕ*_*M*_),, and 1_[·]_ as the indication function that when [·] meets, 1_[·]_ = 1, otherwise, 1_[·]_ = 0. Formally, we have the joint likelihood function as:

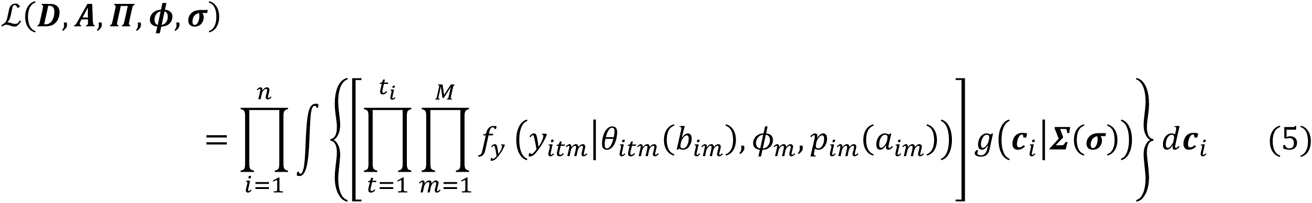

where *f*_*y*_ is the conditional probability density function and given as

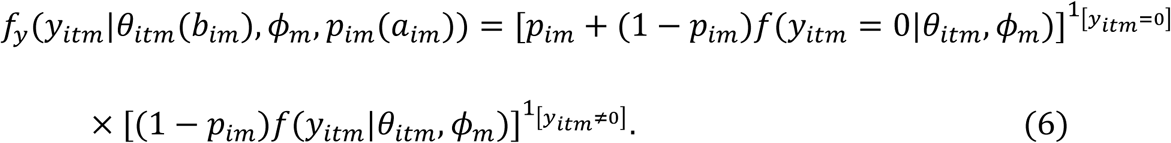

The function *g*(***c***_*i*_|***∑***(***σ***)) is the joint distribution of ***c***_***i***_, and 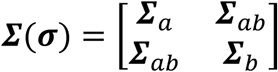 represents the corresponding 2*M* × 2*M* covariance matrix, where ***σ*** accounts for all unique non-zero elements of ***∑***. For the model and computational simplicity, we assume *Cov*(***a***_*i*_, ***b***_*i*_) = ***∑***_*ab*_ = 0, i.e. ***a***_***i***_ and ***b***_*i*_ are independent.

Assuming that the true underlying fixed effects ***A*** and ***D*** are sparse, we advocate a Lasso-type approach, which adds an *ℓ*_1_-penalty for the fixed-effects to the likelihood function. Thus, we consider the following objective function:

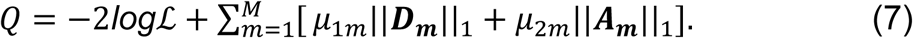

Maximization of the penalized log-likelihood function corresponding to equation (7) with respect to (***D, A, Π, ϕ, σ***) is a computationally challenging task. This is mainly because both integrals with respect to the random effects and the zero-inflated structure do not have analytical solutions. Following the conventional methods, we propose to implement a Laplace approximation on the integral of random effects in equation (7) and use the Expectation-Maximization (EM) algorithm to calculate the expectation and compute parameters iteratively, in which the label of zero is treated as “missing data”. The tuning parameters are selected using Bayesian information criterion (BIC).

### 2.2 Stability Properties

The existence of a stationary distribution has been investigated for the log-linear Poisson auto-regression model based on the perturbation technique [36]. Here, we prove the existence of a stationary distribution of a zero-inflated Poisson mixed-effect auto-regression model in Theorem 1 utilizing the theory of Markov chains which has been proposed to prove the existence of a stationary distribution of a general class of time series count models [37]. The detailed proof is provided in the Supplementary Material, Section 3.

**Theorem 1**. Assuming that time-independent parameters ***η***_*i*_ and ***p***_*i*_ are known, if all eigenvalues of matrix ***B*** lie inside the unit circle, a strict-sense stationary ergodic process {***Y***_*it*_}_*t*∈**N**_ will exist, where **N** denotes the set of natural numbers.

With this Theorem, we can first show that for a microbial community, its dynamic process {***Y***_*it*_}_*t*∈**N**_ has a stationary distribution by proving that all eigenvalues of matrix ***B*** lie inside the unit circle. Then, following Ives et al. [30], we consider the return rate and reactivity as two stability measures based on the variability of the stationary distribution for MAR (1) model. Specifically, return rate depends on the rate at which the perturbed microbial community approaches the stationary distribution and reactivity, assesses how strongly population-level microbiome abundances are pulled towards the mean of the stationary distribution. Both are bounded by the largest eigenvalue of ***B***, denoted by max(***λ***_B_). In general, a smaller max(***λ***_B_) indicates the perturbed microbial community approaches its stationary distribution faster, or a system is less reactive, then the microbial community is more stable. The detailed proof is deferred in the Supplementary Material, Section 3.2.

Based on the theory in Ives et al. [30], for a community with multiple species, the covariance matrix of the stationary distribution depends on the covariance matrix of the process error and the interactions between species captured in the matrix ***B***. As illustrated in our Figure 1, when the external perturbation(blue arrow) acts on the community, the ball(microbial community) sitting in a deep bowl in state 2 which represents a relatively stable system, will return to its stationary state faster than the ball sitting in a shallow bowl in state 1 which represents a less stable system. In a stable system, the variance of stationary distribution is only slightly greater than the variance of process error and the variance of species interaction is small. In contrast, in a less stable system, the species interaction will amplify the environmental variance and create large variance in the stationary distribution, therefore the variance of species interaction is large, assuming the process errors are similar in the compared two states. Thus, the difference between the variances of stationary distribution of different communities can be attributed to species interactions. The smaller of the variance of matrix ***B***, the more stable of the study microbial community.

## 3. Results

### 3.1. Simulation Study

We have conducted extensive simulation studies to evaluate the performance of ARZIMM in both model fitting and variable selection by comparing it with the competing methods: penalized Poisson auto-regression (Poisson), penalized log-normal multivariate auto-regression (MAR), and extended generalized Lotka-Volterra (gLV) equations using Bayesian algorithm (MDSINE) [38]. The brief descriptions of these methods are provided in the Supplementary Material: Section 2.

### 3.2. Simulation Design

We generated the longitudinal absolute abundances from zero-inflated Poisson distribution with parameters *p*_*im*_ and *θ*_*imt*_ for each taxon. Since our focus is on the estimation of the interaction matrix ***B***, which depends on the non-zero part, we adopted a simple simulation design for the zero inflation proportions ***p***_*i*_ = (*p*_*i*1_, …, *p*_*iM*_)′. We ignored the individual variations in ***p***_*i*_ by dropping the random effect term ***a***_*i*_ in equation (2). With model *logit*(***p***_*i*_) = ***AW***_*i*_ and by controlling the values of ***W*** and ***A*** respectively, we set the zero inflation proportions ***p***_*i*_ for 20 taxa to mimic the observed sparsity in real data as

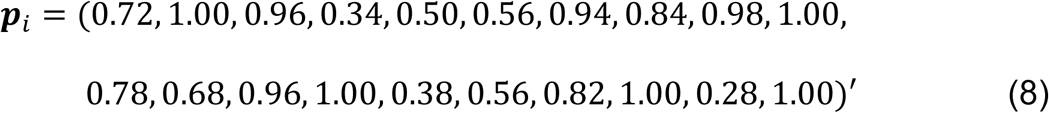

The detailed values of ***W*** and ***A*** are provided in the Supplementary Material, Section 4. We generated the non-zero absolute abundances from Poisson distribution with their ***θ***_*it*_ = (*θ*_*i*1*t*_, …, *θ*_*iMt*_)defined as 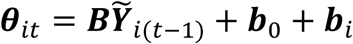, where the intercept ***b***_O_ was set to be the mean log-transformed non-zero absolute abundances of taxa in MIME real data, and the random effects ***b***_*i*_∼*𝒩*(**0**, *diag*(***∑***_b_)) with *diag*(***∑***_b_)∼10^*𝒩*(-1.5,O.5)^. We assumed that the interaction matrix ***B*** was sparse by randomly selecting 5% of its elements to be non-zero. Three interaction matrices were considered with varied informative absolute effect strengths: high 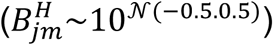, medium 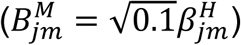, and low 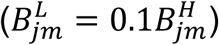, for the non-zero elements *B*_*jm*_. In addition, we designed four simulation scenarios: Scenario 1 with *diag*(***∑***_b_) = **0** and ***p***_*i*_ = **0**, considered as the benchmark situation where subjects are homogeneous and taxa are all presented; Scenario 2 with *diag*(***∑***_b_)∼10^*𝒩*(-1.5.O.5)^ and ***p***_*i*_ = **0**, where subjects are heterogeneous and taxa are all presented; Scenario 3 with *diag*(***∑***_b_) = **0** and ***p***_*i*_ as in (8), where subjects are homogeneous and taxa have zero inflated structure; and Scenario 4 with *diag*(***∑***_b_)∼10^*𝒩*(-1.5.O.5)^ and ***p***_*i*_ as in (8), where subjects are heterogeneous and taxa have zero inflated structure.

In each scenario, we generated 500 independent repetitions for *n* = 20 or 50 subjects, *T* = 10 or 20 time points, and *M* = 20 taxa for each sample to evaluate the performance of ARZIMM.

### 3.1.2. Simulation Results

We first compared the model fittings of ARZIMM, Poisson, and MAR methods using mean normalized squared error score (MNSES), as suggested in the prior studies [39-42]. MNSES is defined as 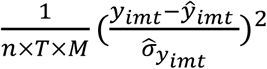 with *ŷ* being the estimated *y*_*imt*_ and 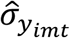 being the estimated standard error of *y*_*imt*_. The closer the MNSES is to 1, the better model fitting the method has. Since MDSINE only provides the estimates of interactions among species without their variance estimates, it was excluded from this comparison. Table 1 and Supplementary Table S1 summarize the median and interquartile range (IQR) of MNSES over 500 replications for these three methods. Overall, the medians of MNSES for ARZIMM are all around the expected value of 1 in various settings across four scenarios, which indicates the good fitness and robustness of ARZIMM in dealing with excess zeros and the correlation among repeated measures at the same time, as well as its satisfying estimation accuracy on the microbial interaction parameters. However, the other two methods: Poisson and MAR, both exhibit inferior performance. The Poisson model is only competent in Scenario 1, when subjects are homogeneous and no excess zeros are present. In Scenarios 2-4, when any factor, excess zero or subject heterogeneity, presents, the predicted values based on the Poisson model deviate greatly from the observed values. Comparing the considered two factors, Poisson model is more sensitive to the subject heterogeneity and presents larger deviations with it. Due to the invalid normality assumption and lack of consideration of the correlation among the longitudinal measurements, the MAR model exhibits the worst performance among three methods with enormous deviation especially in Scenarios 3 and 4, which confirms the inappropriateness of using conventional statistical methods which require the normality assumption to analyze the microbiome data.

**Table 1:**
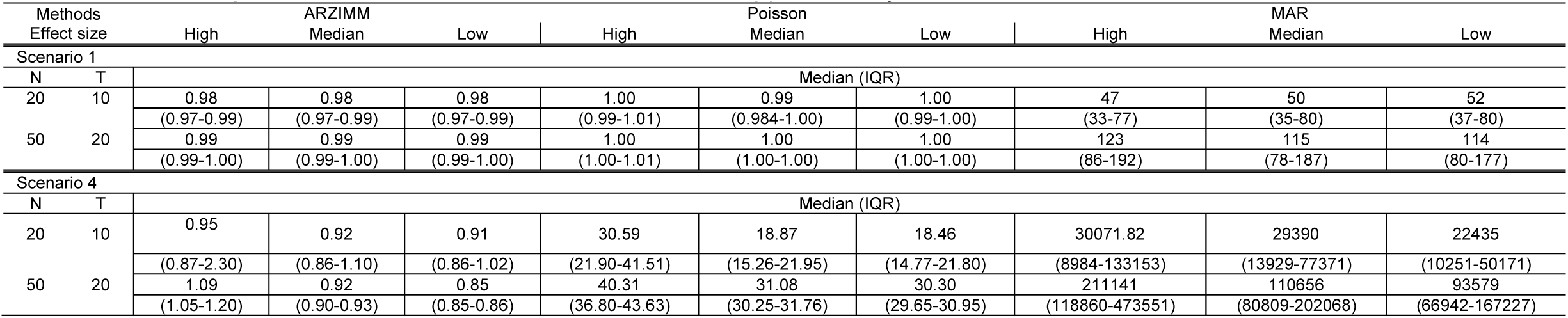
Simulation results for all settings under scenario 1 and 4. Poisson refers to the penalized Poisson autoregression model and MAR refers to penalized log-normal multivariate autoregression model. The reported value is median (IQR) of mean normalized squared error score (MNSES) calculated over 500 simulations for each setting. *N* refers to the number of subjects, and *T* refers to the number of time points. Scenarios 2 and 3 are deferred to Supplementary Material.

Next, we evaluated the variable selection performance for ARZIMM, Poisson, MAR, and MDSINE in terms of true positive rate (TPR; mathematically equals to the power) and false positive rate (FPR; mathematically equals to the type I error). Specifically, TPR quantifies the probability of a significant interaction identified by one method given that the interaction effect is truly nonzero; and FPR quantifies the probability of a significant interaction identified by one method given that the interaction effect is truly zero. The simulation results for 50 subjects with 20 time points are summarized in Figure 3 and all the other simulation results with different subject numbers and time points are deferred to Supplementary Figure S1, because they have a similar pattern as seen in Figure 3. Figure 3 shows that the FPRs of ARZIMM are all at or below the nominal level (5%) across different simulation regimes and effect sizes, and its TPR estimates exhibit a sensible and consistent pattern as they increase as the interaction effect gets stronger across four scenarios. As expected, the FPR and TRP estimates of Poisson and ARZIMM models are coincident under Scenario 1, because when subjects are homogeneous and taxa don’t have excess zeros, ARZIMM model is reduced to Poisson model. However, in Scenarios 2-4, because simple Poisson model fails to take care of the excess zeros or subject heterogeneity, it suffers from the inflated false positives, while ARZIMM does not. For the other two methods, both MAR and MDSINE perform poorly on controlling false positive rates for all simulation scenarios, because MAR fails to fit the skewed and highly sparse microbiome data, while MDSINE captures the interactions based on the averaged effect over subjects in a group but completely ignores the randomness at the subject level process which is the essential characteristic of any biological system. In summary, ARZIMM outperforms the other competitors in handling the excess zeros and subject heterogeneity well with controlled FPR and satisfactory TPR.

**Figure 3:**
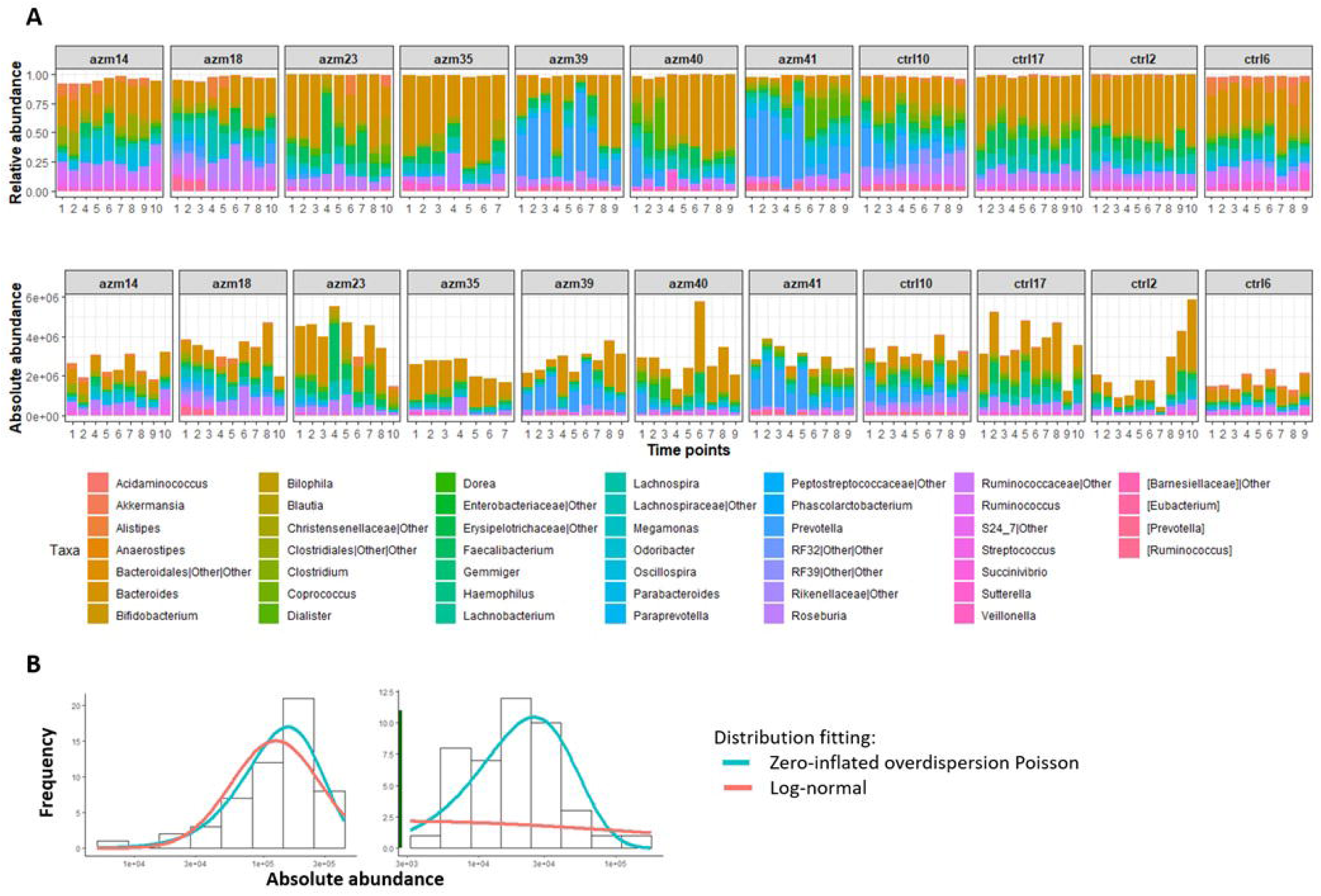
Simulation results of variable selection performance. Poisson refers to the penalized Poisson auto-regression model and MAR refers to penalized log-normal multivariate auto-regression model. MDSINE refers to the method with extended generalized Lotka-Volterra (gLV) equations using a Bayesian algorithm. Mean (and 95% confidence interval) of false positive and true positive rates are reported for 500 simulations with 50 subjects and 20 time points in four scenario: (A) no zero-inflated structure and no heterogeneity, (B) heterogeneity but no zero-inflated structure, (C) zero-inflated structure but no heterogeneity, and (D) both zero-inflated structure and heterogeneity.

To further investigate the performance of informative interaction selection, we calculate Matthew correlation coefficient (MCC), defined as 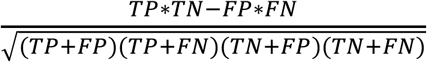, and F-score, defined as 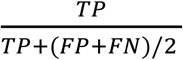, where TP gives the number of selected interactions being true positive, FP gives the number of selected interactions being false positive, TN gives the number of unselected interactions being true negative, and FN gives the number of selected interactions being false negative. MCC ranges from −1 to 1, where value 1 indicates perfect agreement between truth and selection, value −1 indicates perfect disagreement, and value 0 indicates that the selection is random with respect to the truth. F-score ranges from 0 to 1, where value 1 indicates that there are neither false negatives nor false positives and value 0 only indicates no true positives are reported. As expected, MCC and F score are comparable to each other and increase as effect size increases (Supplementary Figure S2). This consistent pattern is observed across four scenarios for ARZIMM but not for Poisson nor MAR models. Similar to TPR and FPR estimates, the MCC and F score values of Poisson and ARZIMM models are coincident under Scenario 1. However, in other situations, both Poisson and MAR perform poorly with low MCC and F score values.

As for the computational cost, ARIZMM took about 2.4 hour to complete the estimation and bootstrap inference for a simulated dataset with 50 subjects, 20 timepoints, and 20 taxa.

### 3.2 Real Data Application

We applied ARZIMM methods to the MIME study. The MIME study is an ongoing randomized trial on 80 healthy volunteers with one control group (ctrl) and two antibiotic groups (amoxicillin, amx, and azithromycin, azm); antibiotics are provided for a 1-week period at the start of the trial. The main microbiome research goal of the MIME study is to evaluate the effects of antibiotics on microbial profiles at both the community and taxonomical levels. With ARIZMM, we propose a different perspective to evaluate the effect of antibiotics through the investigation of microbial interaction and community stability across groups. Because the clinical trial is still ongoing and only partial data are available, the following data analysis is done on a subset of MIME data including only 11 subjects who were randomized to two groups: 4 ctrls and 7 azms. The main purpose of this analysis is to illustrate how to use ARIZMM, not for the scientific conclusion. For each subject, we collected two baseline microbiome samples, three samples during the course of antibiotics, and five post-antibiotic samples. The gut microbiota of these individuals were profiled using 16S rRNA gene targeted sequencing on the Illumina MiSeq platform. To obtain the microbial absolute abundances, we multiplied the relative abundances of OTUs by the sample density 1.1g/cm^3^ and the number of universal 16S rRNA per gram measured using qPCR [43]. In our analysis, samples that collected before treatment in both antibiotic groups were excluded. The abundances of taxa were agglomerated at the genus level and taxa were further filtered if (1) the average relative abundances over all samples are less than 0.1%, and (2) the taxa are presented in less than 5 samples within each group.

First, Figure 4A shows a comparison of the relative abundance (top panel) and the absolute abundances determined by quantitative sequencing (bottom panel) of the dominant bacterial genera in 99 fecal samples from 11 subjects (blocks) across seven to nine time points (shown from left to right within each block) of this preliminary dataset. It is evident that the relative abundance and absolute abundance data present different information about the microbial profiles, and that the total bacterial load changes over time for each subject (i.e., within each block). Thus it is essential to study the microbial interactions using the absolute abundance data.

**Figure 4:**
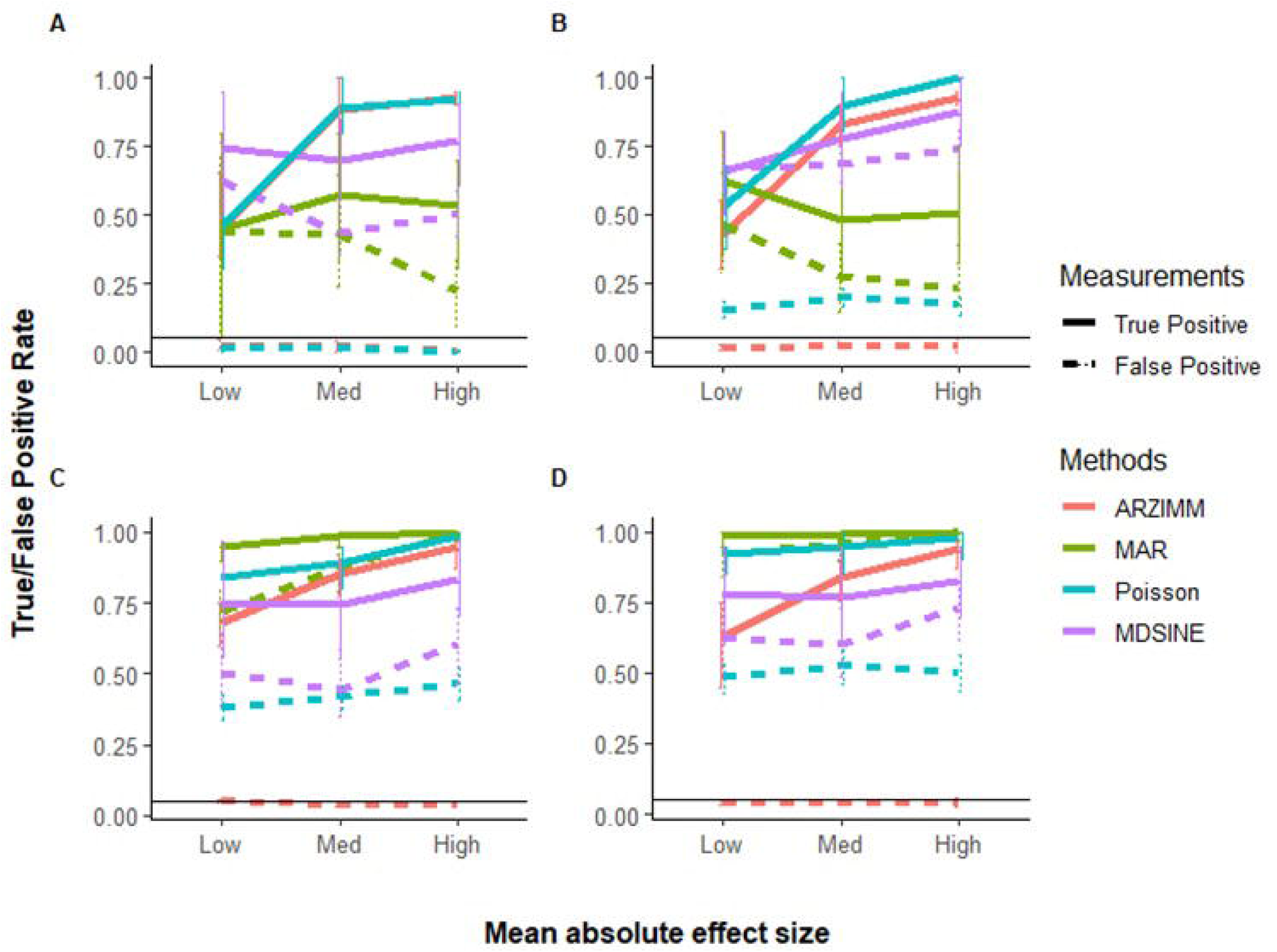
MIME study microbiome data. (A) Difference between relative abundances (top panel) and absolute abundances based on qPCR (bottom panel) of dominant genera in XX fecal samples obtained from 21 subjects (block) at 7–9 time points (x-axis) each. (B) Distribution of absolute abundances of two representative genera from the MIME study, shown in the left and right panels, respectively. For each genus, the absolute abundance is fitted with a log-normal distribution (red line) or a two-part distribution: a zero part (dark green line shown in right panel) and a non-zero part fitted with an over-dispersion Poisson distribution (blue line).

Then, we evaluate the model fitting of the log-normal distribution (used in MAR(1)) and zero-inflated over-dispersed Poisson distribution (used in ARZIMM) on the available subset of MIME data using chi-square goodness of fit test at 5% significance level taxon by taxon. Out of 45 taxa in the control group, 1 and 44 of their absolute abundances were fitted well (*p*>0.05) by log-normal distribution and zero-inflated over-dispersed Poisson distribution respectively. The log-normal distribution fails to fit the data well when microbial taxa’s absolute abundance data are left-skewed and sparse (two examples are illustrated in Figure 4B).

Next we demonstrate how to conduct inference for microbial interactions and community stability with ARZIMM on MIME data. First, we fit ARZIMM to ctrl and azm groups separately, adjusting for age, gender, and BMI, to get their estimated interaction matrix 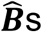. Table 2 reports the characteristics of microbial interaction matrix estimates 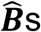. Defining the interaction effect as informative if its 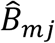’s 95% bootstrap confidence interval (based on 100 bootstrap samples) does not contain zero, we identified 125 and 105 informative interactions, respectively, in azm and ctrl groups. Their interaction effects are illustrated using networks in Figure 5. With more informative interactions, the azm groups have bigger and more complex networks than the ctrl group (first row of Figure 5), while the control group has more large estimated interaction effects than those in azm group as showed in Table 2 and the last three rows of Figure 5. This observation indicates that the antibiotic treatment reduce the strength of the interactions among the taxa and create more variations with more weak interactions among taxa, thus reduce its stability. In the last row of Table 2, based on our stability theory we report the stability properties of the studied microbial communities. The ctrl group has the lower estimates of maximum eigenvalue squared 0.11 comparing to the azm group’s maximum eigenvalue squared 0.32, which indicates that the control microbial community is more stable than the antibiotic communities.

**Table 2:**
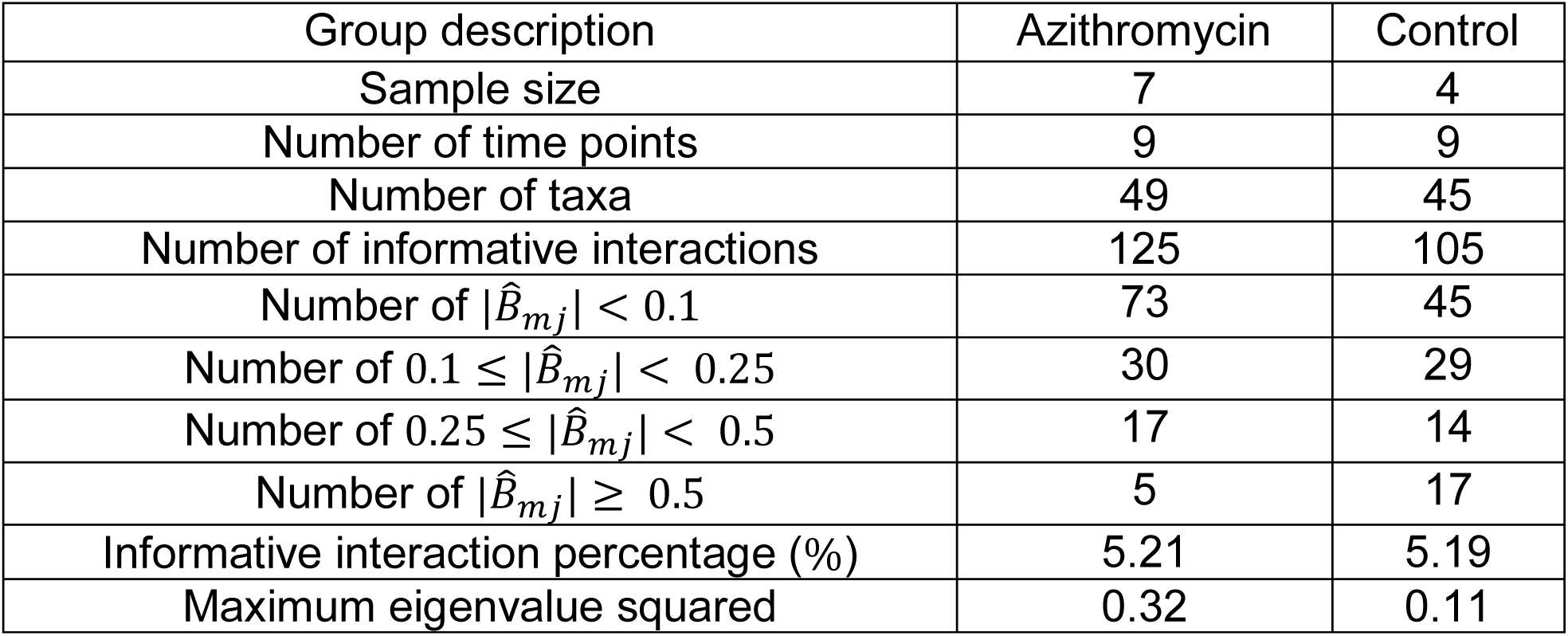
The characteristics of networks.

**Figure 5:**
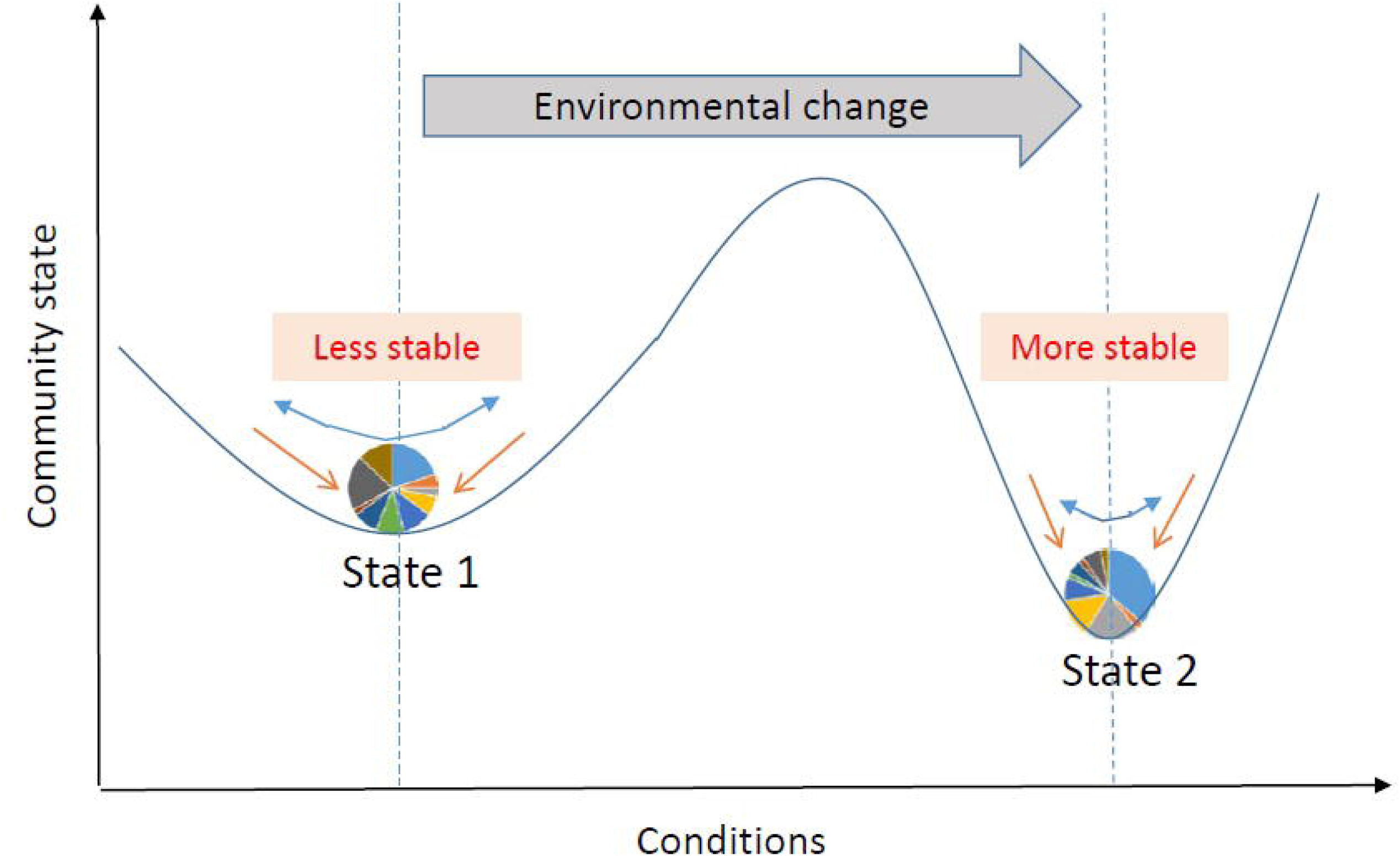
Interaction network. Estimated interaction network for: (A) azithromycin (azm), and (B) control groups, displaying (1) all selected interactions, (2) interactions with 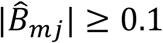, (3) interactions with 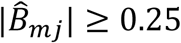, and (4) interactions with 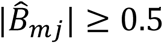. Each node represents a taxon at the genus level, the size of which shows the degree of that taxa and the color of which shows the phylogenetic Order level for each taxon. Each edge with arrow represents an interaction effect, the width of which represents the absolute effect size on a log10 scale, with the color showing a positive (orange) or negative (blue) effect.

Figure 6 provides additional information on the network feature comparison between ctrl and azm groups. Figure6A displays the distribution of the positive and negative informative interaction estimates separately. The ratios between the numbers of positive and negative interactions are both around 1:1 in two groups. Figure 6B presents the frequency distribution of vertex degree of all the taxa in each group and they are all skew to the right. In the figure, a vertex represents a taxon in a community and its vertex degree is the number of informative interaction effect it has with the other taxa. By defining average neighbor degree as the average number of a given taxon’s neighbor vertices’ degrees, Figure 6C shows that the average neighbor degree is negatively correlated with the vertex degree in azm antibiotic treated group, but not in the control group. This indicates that there may be a group of taxa interacting with each other actively in the antibiotic group. It would be interesting to identify such sub-community with additional effort.

**Figure 6:**
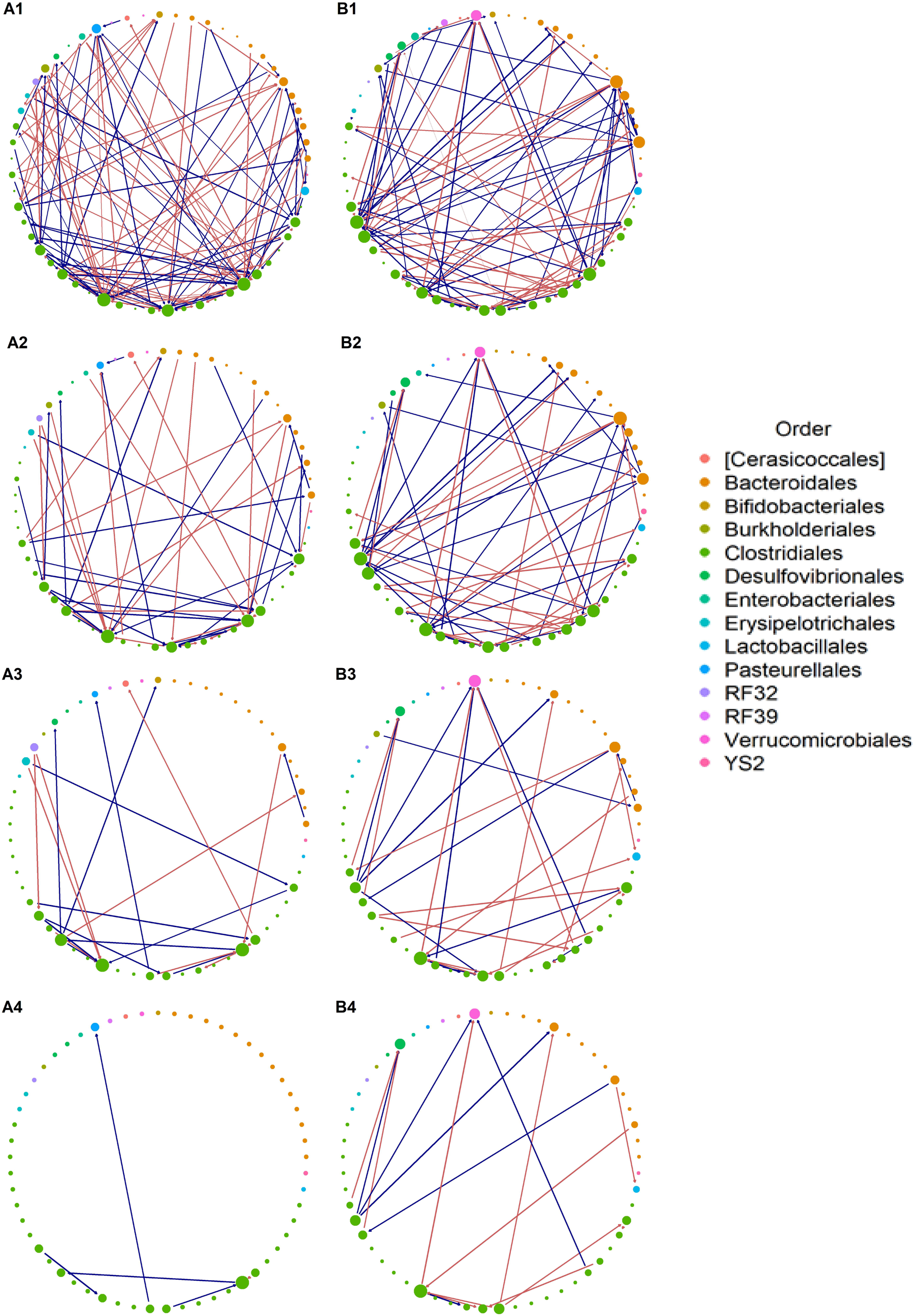
Characteristics of estimated interactions. (A) The effect size of estimated informative interactions, wherein the x-axis represents the log10 scaled absolute effect size, the y-axis represents the count of informative interactions, and the colors represent the positive or negative effects. (B) Histogram of vertex degree, wherein given a vertex, vertex degree is defined as the counts of edges upon the vertex. (C) The average neighbor degree (y-axis) versus vertex degree on a log-log scale (x-axis). The average neighbor degree is the average number of a given taxon’s neighbor vertices’ degrees. Dotted lines represent 95% confidence limits.

## Discussion

In this paper, we propose ARZIMM, an analytic platform which estimates the microbial interactions and community stability using longitudinal microbiome data. ARZIMM tackles the zero-inflated absolute abundance with a mixture distribution of zero and exponential dispersion distribution family, and enhances statistical efficiency by utilizing a random-effects term to account for the correlations among repeated measurements.

It is well-known that microbial correlations calculated from relative abundances are distorted by the compositional nature of microbiome data, and are insufficient in tracking microbial dynamics[44]. We advocate to investigate the microbial correlations using longitudinal absolute abundances which can be determined by combining gene amplicon sequencing with auxiliary total DNA quantitation data. qPCR is one of the most commonly used strategies to quantify total DNA[45] and has been implemented in various statistical analyses[46, 47]. Other alternative methods to quantify the absolute abundances include the combination of the sequencing approach (16S rRNA gene) with robust single-cell enumeration technologies (flow cytometry)[48] and the usage of synthetic chimeric DNA spikes[49].

Plenty of zero-inflated mixed effects models have been recently proposed to handle the excess zeros in microbiome abundance data such as zero-inflated Poisson, negative binomial and quasi-Poisson models[50, 51]. However, none of the existing methods estimates the microbial interactions and community stability. To fill this gap, we extended a zero-inflated Poisson model with auto-regression and random effects modeling, which plays crucial role in efficiently handling the individual heterogeneity and enable the investigation of microbial interactions.

We investigated two community stability measurements derived from ARZIMM: the return rate and reactivity, to further understand ecological dynamics. The estimated interaction matrix ***B*** from the ARZIMM model serves the basis to calculate the largest eigenvalue of ***B***: max(*λ*_***B***_), which determines the return rate of the mean of the transition distribution from the departure to the mean of the stationary distribution. We proposed to measure the reactivity of a microbial community by the expected change of the stationary distribution’s mean in distance from one time point to the next time point. In ARZIMM, higher reactivity coincides with larger eigenvalues of ***B***, thus governed again by max(*λ*_***B***_). Other measures of community stability, such as variance of the stationary distribution [30], warrant further investigations.

It is worth noting that by utilizing the ARZIMM model framework, the time-dependent perturbation (for instance, diet) can also be assessed flexibly in both the autoregressive part and the logistic part in the model. However, the stability based on the microbial interactions has to be interpreted with caution, since the mean of stationary distribution changes along with the time-dependent covariates.

We have demonstrated that ARZIMM outperforms the competing methods and exhibits its feasibility for examining microbial interactions and stability based on longitudinal microbial data. We applied our method to a real human microbiome study of antibiotic treatment and elucidated the microbial interaction network of bacteria from antibiotic and non-antibiotic groups separately. The application of ARZIMM to temporal microbiome data shows great promise. Still, the development of accurate predictive models will require further developments. For example, the method used here to infer microbial interactions may be expanded by adding functional information as well as phylogenetic information. Although this method is primarily developed for the gut microbiota, it may be potentially applied to longitudinal data from any ecological systems. Since interactions between members of microbial communities are primary driving forces for the long-term stability[52], the corresponding stability properties will provide useful principles for community dynamics.

Note that the proposed ARIZMM assumes the probability of observing a zero count for a taxon is constant over time. The reason is two-fold. 1) One major goal of ARIZMM is to derive the inference on the stability of the microbial community over a certain period. With the constant probability of observing a zero count assumption, the stability inference will solely depend on the estimation of the taxon-by-taxon interaction matrix *B*. Otherwise, a stationary distribution won’t exit. 2) Using the MIME data, we estimated the proportions of zeros (denoted as *q*_*mt*_) for all taxa by group at all time points, then calculated the mean 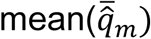 and standard deviation 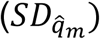 over all the time points and the coefficient of variation 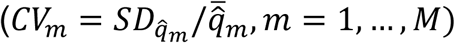 to evaluate their temporal variations. The median of *CV*_*m*_ over all taxa in the control, Amoxicillian and Azithromycin groups are 0.16, 0.12, and 0.34 respectively. This results reveal two observations: 1) the temporal variations of *q*_*mt*_ in most taxa are relative weak; and 2) the temporal variation of the proportions of zero is heterogeneous and there may be no one perfect model fitting all the taxa well. Thus, we believe our assumption that *p*_*m*_ is constant over time is valid and pragmatic. To further check the robustness of our proposed model, we conducted additional simulation by introducing extra randomness when we generate the probability of observing a zero count across the time points, while analyze the data using our proposed model. Our results show that the moderate temporal variation in probability of zero count does not affect ARIZMM’s performance much in capturing the informative interactions by estimating *B* when the absolute effect strengths of interaction matrices is high(FDR<0.05) or medium (FDR <0.15). The detailed simulation design and results are reported in the Supplementary Material Section 4.2 and Figure S3.

The proposed method, ARZIMM has a few limitations and future works are needed to improve it. ARZIMM adopts a simple correlation structure that the random effects in the multivariate logistic component and the multivariate autoregressive component ***a***_***i***_ and ***b***_*i*_ are assumed independent. We took this parsimonious model based on our experience[53, 54] in modeling the longitudinal microbiome data to ease the computational burden. The more general random effects structure with cross-part correlations can provide more robust modeling, however, can suffer from model convergence as well. Further investigation is warranted.

## AUTHORS CONTRIBUTIONS

LH and HL developed the methodological ideas. LH implemented the methods, performed the simulations and real data analysis, and developed the software package. CW and JH contributed to the simulation design and real data analysis. ZG, EF, SH, MJB contributed to the acquisition of utilized real microbiome data. MJB provided biological insights and interpretation of the real data analysis. LH and HL wrote the manuscript. All authors read, edited, and approved the final manuscript.

## DATA AVAILABILITY STATEMENT

Software for implementing the method described in this paper is publicly available on GitHub at https: https://github.com/Hlch1992/ARZIMM.

