## supplementary for "ARZIMM: A Novel Analytic Platform for the Inference of Microbial Interactions and Community Stability from Longitudinal Microbiome Study"

### Supplementary Material

#### 1. Computational algorithm of ARZIMM

Maximization of the penalized log-likelihood function corresponding to equation (2) with respect to $(\boldsymbol{D}, \mathbf{A}, \boldsymbol{\Pi}, \boldsymbol{\phi}, \boldsymbol{\sigma})$ is a computationally challenging task. This is mainly because both the integral with respect to the random effects and the zero-inflated structure do not have analytical solutions.

The Expectation-Maximization (EM) algorithm [1] has been traditionally used under a zero-inflated model in which the labels of zeros are treated as "missing data". Define the Bernoulli random variable $U_{imt} = 1$ if $y_{imt}$ is drawn from zero state, $U_{imt} = 0$ if $y_{imt}$ is drawn from distribution $F$. Thus the "complete" data for the EM algorithm are $(\boldsymbol{y}, \boldsymbol{u})$. Here, $(\boldsymbol{u})$ play the role of missing data, denoted as $\boldsymbol{u}_{it} = (u_{i1t}, \ldots, u_{iMt})'$. Based on $(\boldsymbol{y}, \boldsymbol{u})$, the complete data likelihood is given by

$$\mathcal{L}^c = \prod_{i=1}^{N} \int \{[\prod_{t=1}^{t_i} \prod_{m=1}^{M} f_y^c(y_{imt}, u_{imt} | \theta_{imt}, \phi_m, p_{im})] g(\mathbf{c}_i | \boldsymbol{\Sigma})\} d\mathbf{c}_i$$

where $f_y^c(y_{imt}, u_{imt} | \theta_{imt}, \phi_m, p_{im}) = f(y_{imt} | \theta_{imt}, \phi_m)^{(1-u_{imt})} f_u(u_{imt} | p_{im})$.

Laplace approximations are widely used for maximum likelihood estimation of parameters involving the integral of random effects. Generally, the Laplace approximation is used to calculate integrals of the form.

$$\mathcal{L}^c = \prod_{i=1}^{N} \int e^{-S_i(\mathbf{c}_i)} d\mathbf{c}_i$$

where $S_i(\mathbf{c}_i)$, can be simply defined as $-\log\{[\prod_{t=1}^{t_i} \prod_{m=1}^{M} f_x^c(y_{imt}, u_{imt} | \theta_{imt}, \phi_m, p_{im})] g(\mathbf{c}_i | \boldsymbol{\Sigma})\}$ in our optimization problem, is a known function of variable $\mathbf{c}_i$. Let $\tilde{\mathbf{c}}_i = argmin_{\mathbf{c}_i} S_i(\mathbf{c}_i)$, then the Laplace approximation is given by $\mathcal{L}_i^{c,LA} = \sqrt{2\pi} |S_{i''}(\tilde{\mathbf{c}}_i)|^{-1/2} e^{-S_{i''}(\tilde{\mathbf{c}}_i)}$ where $S_{i''}(\tilde{\mathbf{c}}_i) = \frac{\partial^2 S_i(\mathbf{c}_i)}{\partial \mathbf{c}_i^2} |_{\mathbf{c}_i = \tilde{\mathbf{c}}_i}$. By implementing a Laplace approximation to (2) that

$$Q^{c,LA} = -2\log\mathcal{L}^{c,LA} + \sum_{m=1}^{M} [\mu_{1m} ||\boldsymbol{D}_m||_1 + \mu_{2m} ||\boldsymbol{A}_m||_1],$$

the estimator can be obtained via EM algorithm that calculate the expectation of $Q^{c,LA}$ and compute parameters iteratively as algorithm 1.

---

**Algorithm 1** computational algorithm for ARZIMM

---

Given $\boldsymbol{D}^{(0)}, \mathbf{A}^{(0)}, \boldsymbol{\Pi}^{(0)}, \boldsymbol{\sigma}^{(0)}, \boldsymbol{\phi}^{(0)}$:

**Repeat** for *h*=0,1,..

E-step (Outer): calculate the expectation of $Q^{c,LA(h+1)}$.

**Repeat** for *s*=0,1,...

Given $\mathbf{u}_{it}^{(h)(0)}(\mathbf{c}_i^{(h)}) = \mathbf{u}_{it}^{(h)}(\mathbf{c}_i^{(h)})$, where $\mathbf{u}_{it}^{(0)} = (1_{\{y_{i1t}=0\}}, \dots, 1_{\{y_{iMt}=0\}})'$:

M-step (Inner): solve the following minimization problem:

$$\mathbf{c}_i^{(h)(s+1)} = \underset{\mathbf{c}_i}{\operatorname{argmin}} S_i(\mathbf{c}_i | \mathbf{u}_{it}^{(h)(s)}).$$

E-step (Inner): estimate $\mathbf{u}_{it}^{(h)(s+1)}(\mathbf{c}_i^{(h)(s+1)})$, that for m-th element

$$u_{it}^{m(h)(s+1)}(\mathbf{c}_i^{(h)(s+1)}) = \begin{cases} [1 + e^{-logi\ (p_{im}(\mathbf{c}_i^{(h)(s+1)})) + \log f(0)}]^{-1} & y_{imt} = 0 \\ 0 & y_{imt} \neq 0 \end{cases}$$

**until** convergence. Calculate the expectation of $Q^{c,LA(h+1)}$.

M-step (Outer): solve optimization problem to obtain the estimate

$$\boldsymbol{D}^{(h+1)}, \mathbf{A}^{(h+1)}, \boldsymbol{\Pi}^{(h+1)}, \boldsymbol{\sigma}^{(h+1)}, \boldsymbol{\phi}^{(\mathrm{h}+1)} = \underset{\boldsymbol{D},\boldsymbol{A},\boldsymbol{\Pi},\boldsymbol{\sigma},\boldsymbol{\phi}}{\operatorname{argmin}} Q^{c,LA(h+1)}$$

**until** convergence.

### 2. Competing methods

#### 2.1 Multivariate autoregression method (MAR)

Multivariate autoregression (MAR(1)) models are applied to estimate the ecological interactions after a log transformation of ecological time series counts[2], formulated as

$$\boldsymbol{X}_t = \boldsymbol{A} + \boldsymbol{B}\boldsymbol{X}_{t-1} + \boldsymbol{E}_t$$

where $\boldsymbol{X}_t$ is a vector of (log-transformed) population abundances at time $t$, $\boldsymbol{A}$ is a vector of constants, $\boldsymbol{B}$ is a matrix whose elements $b_{ij}$ give the effect of the abundance of species $j$ on the per capita population growth rate of species $i$, and $\mathbf{E}_\mathrm{t}$ is a vector of process errors that has a multivariate normal distribution with mean vector $\mathbf{0}$ and covariance matrix $\boldsymbol{\Sigma}$.

We implement the regularized maximum likelihood (ML) estimation procedure to select variables and estimate parameters simultaneously. Assuming that the true underlying effects $\boldsymbol{B}$ are sparse, we advocate a Lasso-type approach, which adds an $l_1$-penalty for the effects to the likelihood function. The tuning parameters are selected using Bayesian information criterion (BIC).

#### 2.2 Poisson auto-regression method (Poisson)[2]

In order to model the longitudinal sparse microbial counts from a study cohort, let $Y_{imt}$ denote the observed count of bacterial taxon $m(m = 1, \dots, M)$ for subject $i$ at time point $t(i =$

$1,2,\dots,N, t = 1,\dots,T_i$). Consider a Poisson distribution of microbial count that $(Y_{imt}+1) \sim Poisson(\theta_{imt})$, where $Poisson$ represents the Poisson distribution with the parameter $\theta$. Considering the longitudinal microbial counts, the Poisson autoregression is performed as:

$$\boldsymbol{\theta}_{it} = \boldsymbol{A} + \boldsymbol{B}\boldsymbol{Y}_{i(t-1)},$$

Where $\boldsymbol{\theta}_{it} = (\theta_{i1t}, \dots, \theta_{iMt})'$ and $\boldsymbol{Y}_{i(t-1)} = \left(Y_{i1(t-1)}, \dots, Y_{iM(t-1)}\right)'$.

Similar regularized ML estimation procedure is implemented by adding an $l_1$-penalty for the effects to the likelihood function. Bayesian information criterion (BIC) is applied for the tuning parameters selection.

#### 2.3 The Microbial Dynamical Systems INference Engine (MDSINE)

To infer dynamical system models of the microbiota from high-throughput time-series data.The Microbial Dynamical Systems INference Engine (MDSINE)[3] software package implements the extended generalized Lotka-Volterra (gLV) equations model for $M$ taxa measured in $N$ subjects, the rate of change of the concentration of taxon $m$ in subject $i$, expressed as:

$$\begin{aligned}\frac{df_{mi}}{dt} &= \alpha f_{mi}(t) + \sum_{j=1}^{M} \beta_{mj} f_{mi}(t) f_{ji}(t) + \sum_{p=1}^{P} \gamma_{mp} f_{mi}(t) u_p(t)\\ &\approx \alpha \hat{f}_{mi} + \sum_{j=1}^{M} \beta_{mj} \hat{f}_{mit} \hat{f}_{jit} + \sum_{p=1}^{P} \gamma_{mp} \hat{f}_{mit} u_p(t)\end{aligned}$$

where $\boldsymbol{\alpha}$ represent unbounded growth rates, the $\boldsymbol{\beta}$ represent pair-wise microbe-microbe interactions, and the $\boldsymbol{\gamma}$ represent effects of $P$ perturbations. The functions $u_p(t)$ are binary valued, indicating if the given perturbation is present at time $t$. $\hat{f}_{mit}$ represents an estimate of the concentration of taxon $m$ in subject $i$ at time point $t$.

The MDSINE software is is publicly available [3]

### 3. Ergodicity and stability measures

#### 3.1 Ergodicity

In this section, we establish the existence and uniqueness of the invariant distribution of the observation process $\{\boldsymbol{Y}_{it}\}$ based on ARIZMM, and shows the ergodicity and existence of some moments for its stationary distribution. The results are formed under the assumption that the time independent parameters $\boldsymbol{\eta}_i$ and $\boldsymbol{p}_i$ are known. To derive the general conditions, we introduce an unobserved Markov process $\{\boldsymbol{X}_{it}\}$ based on a measurable function $y \mapsto f_{\boldsymbol{\eta}_i}(y)$ and a Markov kernel $\mathcal{ZIP}_{\boldsymbol{p}_i}$ from $(\mathbf{R}^M, \mathcal{B}(\mathbf{R}^M))$ to $(\mathbf{N}^M, \mathcal{B}(\mathbf{N}^M))$, which satisfies the following recursions:

$$\begin{aligned}\boldsymbol{Y}_{it} | \mathcal{D}_{i(t-1)} &\sim \mathcal{ZIP}_{\boldsymbol{p}_i}(\boldsymbol{X}_{i(t-1)})\\ \boldsymbol{X}_{it} &= f_{\boldsymbol{\eta}_i}(\boldsymbol{Y}_{it})\end{aligned} \quad \text{(S1)}$$

where $\mathcal{B}(\cdot)$ denotes the Borel $\sigma$-algebra for $(\cdot)$. Specifically, $f_{\boldsymbol{\eta}_i}(\boldsymbol{Y}_{it})$ is defined as $\boldsymbol{\theta}_{it}$ satisfying (3) and $\mathcal{ZIP}_{\boldsymbol{p}_i}(\boldsymbol{\theta})$ is a zero-inflated Poisson distribution satisfying (1) where each dimension $Y^m$ is independently distributed as a zero-inflated Poisson with parameters $(p^m, \theta^m)$. We will give the assumptions so that the unobserved Markov process $\{\boldsymbol{X}_{it}\}$ converge to the stationary distribution. Then, we will show the existence of a strict-sense stationary ergodic process on $\{\boldsymbol{Y}_{it}\}_{t\in\mathbf{N}}$, the solution to the recursion (S1). Finally, we will derive the conditions meeting the assumptions for stationarity.

Based on the theory of Markov chains without irreducibility assumption to prove the existence of a stationary distribution, the existence of a stationary distribution has been investigated for the log-linear Poisson autoregression model[4]. We follow this approach and extend it to a zero-inflated multivariate version.

For a general Markov chain $\boldsymbol{X}_i = \{\boldsymbol{X}_{it}\}_{t\in N}$ on state space $\mathbf{R}^M$ with $\sigma$-algebra $\mathcal{B}(\mathbf{R}^M)$, define $T_i^t(\mathbf{x}_i, A) = Pr(\mathbf{X}_t \in A|\mathbf{X}_{i0} = \mathbf{x}_i)$ for $A \in \mathcal{B}(\mathbf{R}^M)$ to be the $t$-step transition probability starting from state $\mathbf{X}_{i0} = \mathbf{x}_i$. Define for all $(\mathbf{x}_i, \tilde{\mathbf{x}}_i) \in \mathbf{R}^M \times \mathbf{R}^M$, $\bar{T}_i(\mathbf{x}_i, \tilde{\mathbf{x}}_i; \cdot)$ as the law of $(\mathbf{X}_{i1}, \widetilde{\mathbf{X}}_{i1})$ that

$$\mathbf{X}_{i1} = f_{\boldsymbol{\eta}_i}(\mathbf{Y}_{it}); \quad \widetilde{\mathbf{X}}_{i1} = f_{\boldsymbol{\eta}_i}(\mathbf{Y}_{it}); \quad Y_{it} \sim \mathcal{ZIP}_{\boldsymbol{p}_i}(\mathbf{x}_i^{\vee}) \qquad \text{(S2)}$$

where $\boldsymbol{x}_i^{\vee} = (x_i^1 \vee \tilde{x}_i^1, \ldots, x_i^M \vee \tilde{x}_i^M)^T$. Consider the following assumptions:

(A1) The Markov kernel $T_i$ is a Feller semigroup. Moreover, there exist a compact set $C \in \mathcal{B}(\mathbf{N}^M)$, $(b, \epsilon) \in \mathbf{R}_*^+ + \times \mathbf{R}_*^+ +$ and a function $V: \mathbf{R}^M \to \mathbf{R}^+ +$ such that

$$T_i V \leq V - \epsilon + b1_C$$

(A2) The Markov kernel $T_i$ has a reachable point.

(A3) There exist a kernel $\bar{T}_i := \mathrm{Pr}$ on $(\mathbf{R}^M \times \mathbf{R}^M, \mathcal{B}(\mathbf{R}^M)^{\otimes 2})$, a measurable function $\alpha(\boldsymbol{x}_i, \tilde{\boldsymbol{x}}_i) := \exp\{-\sum_{m=1}^{M}(1-p^m)[e^{x_i^m \vee \tilde{x}_i^m} - e^{x_i^m \wedge \tilde{x}_i^m}]\}$, and a measurable function $W(\boldsymbol{x}_i, \widetilde{\boldsymbol{x}}_i) := \sum_{m=1}^{M}(1-p^m)e^{|x_i^m| \vee |\tilde{x}_i^m|}$ such that for all $(\boldsymbol{x}_i, \widetilde{\boldsymbol{x}}_i) \in \mathbf{R}^M \times \mathbf{R}^M$,

$$1 - \alpha(\boldsymbol{x}_i, \widetilde{\boldsymbol{x}}_i) \leq d(\boldsymbol{x}_i, \widetilde{\boldsymbol{x}}_i) W(\boldsymbol{x}_i, \widetilde{\boldsymbol{x}}_i)$$

where $d(\boldsymbol{x}_i, \widetilde{\boldsymbol{x}}_i) = |\boldsymbol{x}_i - \widetilde{\boldsymbol{x}}_i|$ denotes the euclidean distance. There exists $(\rho, \beta) \in (0,1) \times \mathbf{R}$ such that for all $(\mathbf{x}_i, \tilde{\mathbf{x}}_i) \in \mathbf{R}^M \times \mathbf{R}^M$

$$\mathrm{Pr}\{d(\boldsymbol{X}_{i1}, \widetilde{\boldsymbol{X}}_{i1}) \leq \rho d(\boldsymbol{x}_i, \widetilde{\boldsymbol{x}}_i)\} \xrightarrow{a.s.} 1$$

$$\bar{T}_i W \leq W + \beta$$

Moreover, for all $\mathbf{x} \in \mathbf{R}^M$, there exists $\gamma_{\mathbf{x}} > 0$ such that

$$\sup_{\tilde{\mathbf{x}} \in B(\mathbf{x}, \gamma_{\mathbf{x}})} W(\boldsymbol{x}, \widetilde{\boldsymbol{x}}) < \infty$$

where $B(\boldsymbol{x}, \gamma_x)$ is the open ball of radius $\gamma_x$ with respect to $d$ and centered at $\boldsymbol{x}$.

**Theorem 1.** *Assume that all eigenvalues of matrix $\boldsymbol{B}$ lie inside the unit circle. Then, the assumptions (A1)—(A3) hold.*

The proof can be easily adapted from [3].

**Theorem 2.** *Assume that (A1)—(A3) hold. Then, the Markov kernel $T_i$ admits a unique invariant probability measure.*

**Lemma 1.** *Assume that the Markov kernel $T_i$ admits an invariant kernel $\pi$ and that there exist a measurable function $V: \mathbf{R}^p \rightarrow \mathbf{R}^{+}+$ and real numbers $(\lambda, \beta) \in (0,1) \times \mathbf{R}_*^+$ such that $QV \leq \lambda V + \beta$. Then,*

$$\pi V \leq \beta/(1-\lambda) < \infty$$

**Proposition 1.** *Assume that the Markov kernel $T_i$ admits a unique invariant probability measure. Then, there exists a strict-sense stationary ergodic process on $\mathbf{N}$,$\{\boldsymbol{Y}_t\}_{t\in\mathbf{N}}$, the solution to the recursion of the model.*

*Proof.* The property of asymptotically strong Feller Markov chains has been investigated in Hairer et al. [5]. The following theorem is taken from Theorem 3.16 in Hairer et al. [5].

**Definition 1.** *A Markov transition semigroup $T_i$ on a Polish space $\mathbf{R}^p$ is called asymptotically strong Feller at $\boldsymbol{x}$ if there exists a totally separating system of pseudo-metrics $\{d_n\}$ for $\mathbf{R}^p$ and a sequence $t > 0$ such that*

$$\lim_{\gamma\rightarrow\infty} \limsup_{t\rightarrow\infty} \sup_{\tilde{\mathbf{x}}\in B(\mathbf{x},\gamma)} ||T_i^t(\boldsymbol{x},\cdot) - T_i^t(\boldsymbol{x},\cdot)||_{d_n} = 0 \tag{S3}$$

**Theorem 3.** *Assume that $T_i$ is Markov semigroup satisfying (S3) and admits reachable points $\boldsymbol{x}$. Then there exists at most one invariant probability measure for $T_i$.*

#### 3.2 Stability measures

##### Return rate

The first stability measure is the return rate. For a system, where a stationary distribution exists, the more rapid the convergence, the more stable the system. Following Ives et. al [2], we measure the rate of return by the rate at which the transition distribution converges to the stationary distribution from an initial condition $\boldsymbol{Y}_{i0} = \boldsymbol{y}_{i0}$. Previously, we established the existence of the stationary distribution for the observation process $\{\boldsymbol{Y}_{it}\}$ based on an unobserved Markov process $\{\boldsymbol{X}_{it}\}$ with a stationary distribution. Therefore, we consider the return rate of the unobserved Markov process $\{\boldsymbol{X}_{it}\}$. Similar to the MAR(1) process, the return rate of the mean vector of $\boldsymbol{X}_{it}$ is governed by $\boldsymbol{B}$, and $\max(\boldsymbol{\lambda}_B)$ limits the return time of the mean to $\boldsymbol{X}_{i\infty}$.

##### Reactivity

The second stability measure is the reactivity, which assesses how strongly population-level microbiome abundances are pulled towards the mean of the stationary distribution. To quantify the reactivity, Ives et. al [2] considered the expectation over the stationary distribution of the change in distance from the mean of the stationary distribution between successive time steps. Similar to return rate, we consider the reactivity of the unobserved Markov process $\{\boldsymbol{X}_{it}\}$. Let $\Delta\boldsymbol{X}_{it} = \boldsymbol{X}_{it} - \boldsymbol{\mu}_{i\infty}$ be the difference between $\boldsymbol{X}_{it}$ and the mean of the stationary distribution. Therefore, the reactivity is defined as

$$\mathrm{E}_{\boldsymbol{X}_{i(t-1)}}\left[\frac{\mathrm{E}_{\boldsymbol{X}_{it}|\boldsymbol{X}_{i(t-1)}}[||\Delta\boldsymbol{X}_{it}|\Delta\boldsymbol{X}_{i(t-1)}||^2]-||\Delta\boldsymbol{X}_{i(t-1)}||^2}{||\Delta\boldsymbol{X}_{i(t-1)}||^2}\right],$$

where $||\cdot||^2$ denotes the squared Euclidean distance. To obtain the reactivity of the system averaged over a long time period, we can let the distribution of $\boldsymbol{X}_{i(t-1)}$ be the stationary distribution $\boldsymbol{X}_{i\infty}$, and thus, the reactivity can be shown as

$$\frac{\mathrm{E}_{\boldsymbol{X}_{i\infty}}[\Delta\boldsymbol{X}_{i(t-1)}{}'\boldsymbol{B}'\boldsymbol{B}\Delta\boldsymbol{X}_{i(t-1)}]}{\mathrm{E}_{\boldsymbol{X}_{i\infty}}[||\Delta\boldsymbol{X}_{i(t-1)}||^2]}-1\leq\max(\boldsymbol{\lambda}_{\boldsymbol{B}'\boldsymbol{B}})-1,$$

where $\max(\boldsymbol{\lambda}_{\boldsymbol{B}'\boldsymbol{B}})$ is the dominant eigenvalue of the matrix $\boldsymbol{B}'\boldsymbol{B}$. Thus, $\max(\boldsymbol{\lambda}_{\boldsymbol{B}'\boldsymbol{B}})-1$ indicates the "worst-case" reactivity of a system.

### 4. Simulation setting

#### 4.1. Simulation setting for the zero part

We generated the longitudinal absolute counts from zero-inflated Poisson distribution with parameters $p_i^m$ and $\theta_{it}^m$ for each taxon. Since our focus is on the estimation of the interaction matrix $\boldsymbol{B}$, which depends on the non-zero part, we adopted a simple simulation design for the zero inflation proportions $\boldsymbol{p}_i=(p_i^1,\ldots,p_i^M)'$. We ignored the individual variations in $\boldsymbol{p}_i$ by dropping the random effect term $\boldsymbol{a}_i$ in equation (1). In the simulation design, in order to mimic the observed sparsity in real data, we set the zero inflation proportions $\boldsymbol{p}_i$ for 20 taxa as $\boldsymbol{p}_i$= $(0.72, 1.00, 0.96, 0.34, 0.50, 0.56, 0.94, 0.84, 0.98, 1.00, 0.78, 0.68, 0.96, 1.00, 0.38, 0.56, 0.82, 1.00, 0.28, 1.00)'$. In order to test our algorithm on estimating the parameters in zero inflation part along with the non-inflation part, based on the model $logit(\boldsymbol{p}_i)=\boldsymbol{AW}_i$, we constructed six binary covariates $\boldsymbol{W}_i$ with $W_{il}\sim Bernoulli(\omega_l)$; $l=1,2,\ldots,6$, and $(\omega_1,\ldots,\omega_6)^T=(0.05, 0.05, 0.5, 0.5,0.5, 0.5)'$ when $N=20$, and $(\omega_1,\ldots,\omega_6)^T=(0.02, 0.04, 0.1,0.3, 0.5, 0.7)'$ when $N=50$. The elements of $\boldsymbol{\alpha}$ were set to be sparse with informative coefficients to be 5.

#### 4.2. Simulation setting for the mis-specified model

In this section, we conducted simulation experiments to verify the robustness of ARZIMM under model misspecification. Based on Scenario 4, where both zero-inflated structure and heterogeneity are modeled, we added a random term to the zero inflation proportions $\boldsymbol{p}_i$ to allow the zero inflation proportions $\widetilde{\boldsymbol{p}}_i=\boldsymbol{p}_i+\alpha\beta_i$, where $\alpha\sim Bernoulli(0.1)$ and $\beta_i$'s $m$-th element (1) $\beta_{im}\sim N(0, 0.2p_{im})$; or (2) $\beta_{im}\sim N(0, 0.3p_{im})$, to vary along the time. Here, $\alpha$, the strength of variation, was set based on the coefficient of variation estimates using MIME data as we reported in the Discussion. In this mis-specified simulation, we evaluated the variable selection performance for ARZIMM, Poisson, and MAR in terms of true positive rate (TPR) and false positive rate (FPR) and present the results in Supplementary Figure S3. The results show that the moderate temporal variation in probability of zero count does not affect ARIZMM's performance much in capturing the informative interactions by estimating B when the absolute effect strengths of interaction matrices is high(FDR<0.05) or medium (FDR <0.15).

### 5. Supplementary Tables and Figures

**Supplementary Table 1.** Simulation results for all settings under four scenarios. Poisson refers to the penalized Poisson auto-regression model and MAR refers to penalized log-normal multivariate auto-regression model. Reported value are medians (IQRs) of mean normalized squared error score (MNSES) calculated over 500 simulations for each setting.

| Methods | | ARZIMM | | | Poisson | | | MAR | | |
|---|---|---|---|---|---|---|---|---|---|---|
| Effect size | | High | Median | Low | High | Median | Low | High | Median | Low |
| Scenario 1 | | | | | | | | | | |
| *N* | *T* | Median (IQR) | | | | | | | | |
| 20 | 20 | 0.99 (0.98-0.99) | 0.99 (0.98-1.00) | 0.99 (0.99-1.00) | 1.00 (0.99-1.01) | 1.00 (0.99-1.00) | 1.00 (0.99-1.01) | 76 (53-120) | 74 (53-122) | 74 (53-131) |
| 50 | 10 | 0.98 (0.98-0.99) | 0.98 (0.98-0.99) | 0.98 (0.98-0.99) | 1.01 (1.00-1.01) | 1.00 (0.99-1.00) | 1.00 (0.99-1.00) | 78 (55-133) | 77 (55-134) | 84 (59-139) |
| Scenario 2 | | | | | | | | | | |
| *N* | *T* | Median (IQR) | | | | | | | | |
| 20 | 10 | 0.97 | 0.98 | 0.98 | 1.51 | 1.33 | 1.23 | 70 | 56 | 57 |
| | | (0.96-1.48) | (0.96-1.09) | (0.97-0.99) | (1.50-1.53) | (1.32-1.35) | (1.22-1.24) | (49-109) | (40-86) | (38-94) |
| 20 | 20 | 0.99 | 1.06 | 1.00 | 1.49 | 1.34 | 1.25 | 91 | 80 | 79 |
| | | (0.98-1.36) | (0.98-1.08) | (0.99-1.01) | (1.48-1.50) | (1.33-1.35) | (1.23-1.26) | (66-156) | (57-130) | (57-135) |
| 50 | 10 | 0.96 | 1.09 | 1.17 | 1.57 | 1.36 | 1.26 | 89 | 78 | 80 |
| | | (0.96-0.97) | (1.05-1.15) | (1.14-1.21) | (1.56-1.62) | (1.36-1.37) | (1.26-1.27) | (63-148) | (54-124) | (55-127) |
| 50 | 20 | 0.98 | 1.13 | 1.14 | 1.54 | 1.36 | 1.26 | 117 | 116 | 114 |
| | | (0.98-0.98) | (1.07-1.17) | (1.11-1.19) | (1.53-1.56) | (1.36-1.37) | (1.26-1.26) | (82-191) | (79-209) | (79-179) |
| Scenario 3 | | | | | | | | | | |
| *N* | *T* | Median (IQR) | | | | | | | | |
| 20 | 10 | 0.92 | 0.87 | 0.88 | 27.6 | 19.4 | 19.6 | 35012 | 13069 | 9128 |
| | | (0.90-0.96) | (0.85-0.91) | (0.85-0.94) | (22.0-32.8) | (15.4-23.3) | (15.5-23.3) | (10154-101650) | (6552-24509) | (3320-16752) |
| 20 | 20 | 0.94 | 0.92 | 0.92 | 41.3 | 30.0 | 29.4 | 144005 | 32263 | 17706 |
| | | (0.92-1.30) | (0.90-0.94) | (0.90-0.93) | (35.6-45.4) | (26.5-32.5) | (26.4-31.6) | (64379-313924) | (20626-58587) | (10881-27450) |
| 50 | 10 | 0.95 | 0.95 | 0.95 | 38.5 | 28.0 | 26.8 | 131596 | 44419 | 24869 |
| | | (0.94-0.97) | (0.94-0.96) | (0.94-0.96) | (33.4-41.3) | (26.3-29.4) | (25.1-28.1) | (63797-242634) | (27851-83709) | (16184-44889) |
| 50 | 20 | 0.98 | 0.98 | 0.98 | 43.6 | 30.0 | 28.5 | 252042 | 63645 | 39205 |
| | | (0.97-0.98) | (0.97-0.98) | (0.97-0.98) | (42.0-44.8) | (29.3-30.8) | (27.6-29.1) | (164342-395273) | (46459-107196) | (27347-70133) |
| Scenario 4 | | | | | | | | | | |
| *N* | *T* | Median (IQR) | | | | | | | | |
| 20 | 20 | 0.95 | 0.92 | 0.84 | 43.9 | 28.5 | 28.7 | 150843 | 85354 | 43717 |
| | | (0.91-2.32) | (0.89-0.96) | (0.83-0.86) | (32.5-51.4) | (25.6-31.0) | (25.0-30.9) | (55236-446842) | (45584-169844) | (25602-83638) |
| 50 | 10 | 1.32 | 0.87 | 0.84 | 36.5 | 28.3 | 28.0 | 55616 | 67888 | 48800 |

| | | | | | | | | | |
|---|---|---|---|---|---|---|---|---|---|
| | (1.24-1.41) | (0.86-0.89) | (0.83-0.86) | (31.8-42.8) | (26.3-30.0) | (26.2-29.5) | (30732-105802) | (43224-120112) | (33059-89420) |

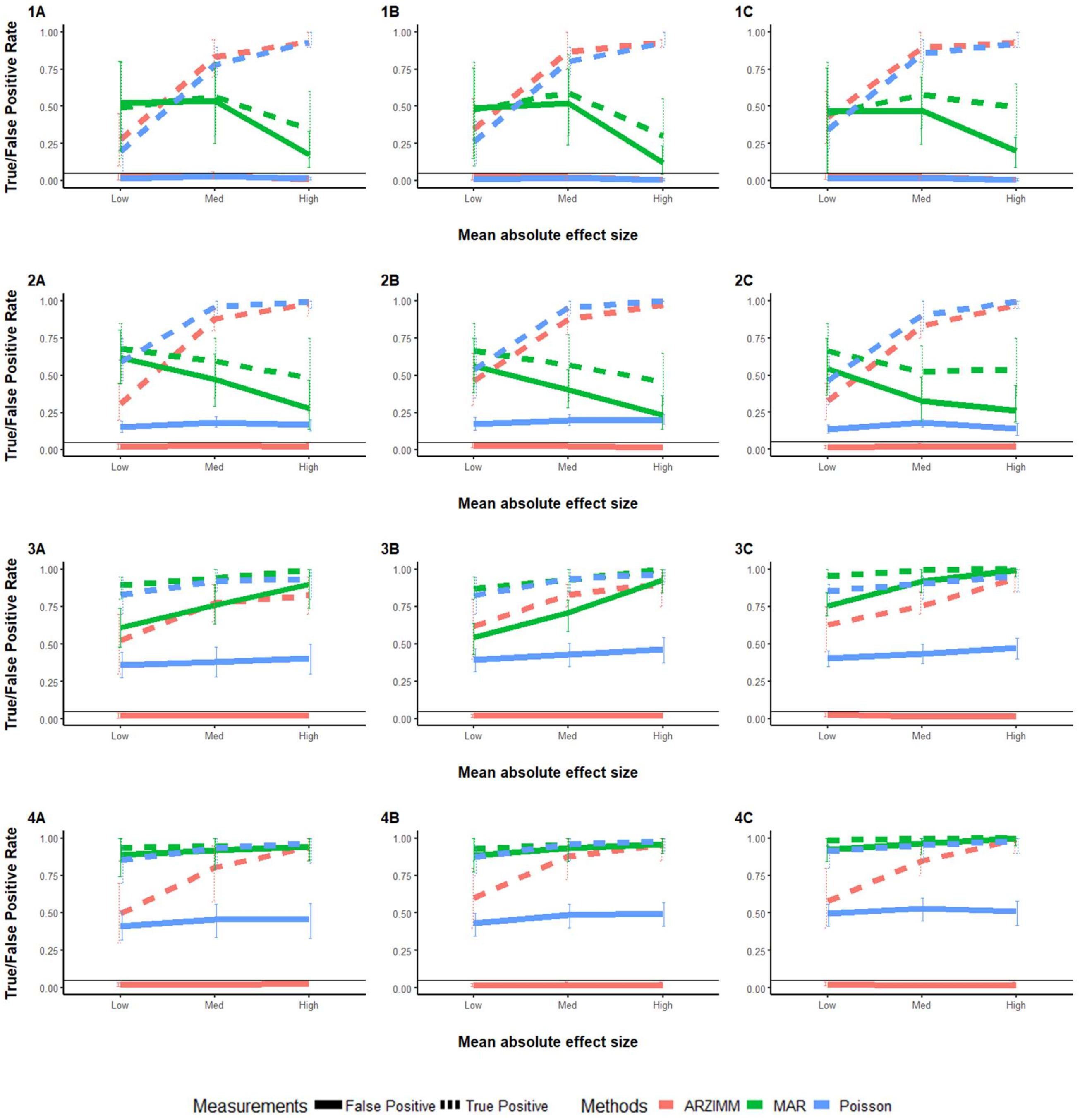


**Supplementary Figure 2.** Simulation results of four scenarios for (A) 20 subjects with 10 time points; (B) 20 subjects with 10 time points; (C) 50 subjects with 10 time points. Poisson refers to the penalized Poisson autoregression model and MAR refers to penalized log-normal multivariate autoregression model. False Positive Rate and True Positive Rate are reported with mean and 95% interval over 500 simulations for four scenarios to have: (1) no zero-inflated structure and no heterogeneity; (2) heterogeneity but no zero-inflated structure; (3) zero-inflated structure but no heterogeneity; (4) both zero-inflated structure and heterogeneity.

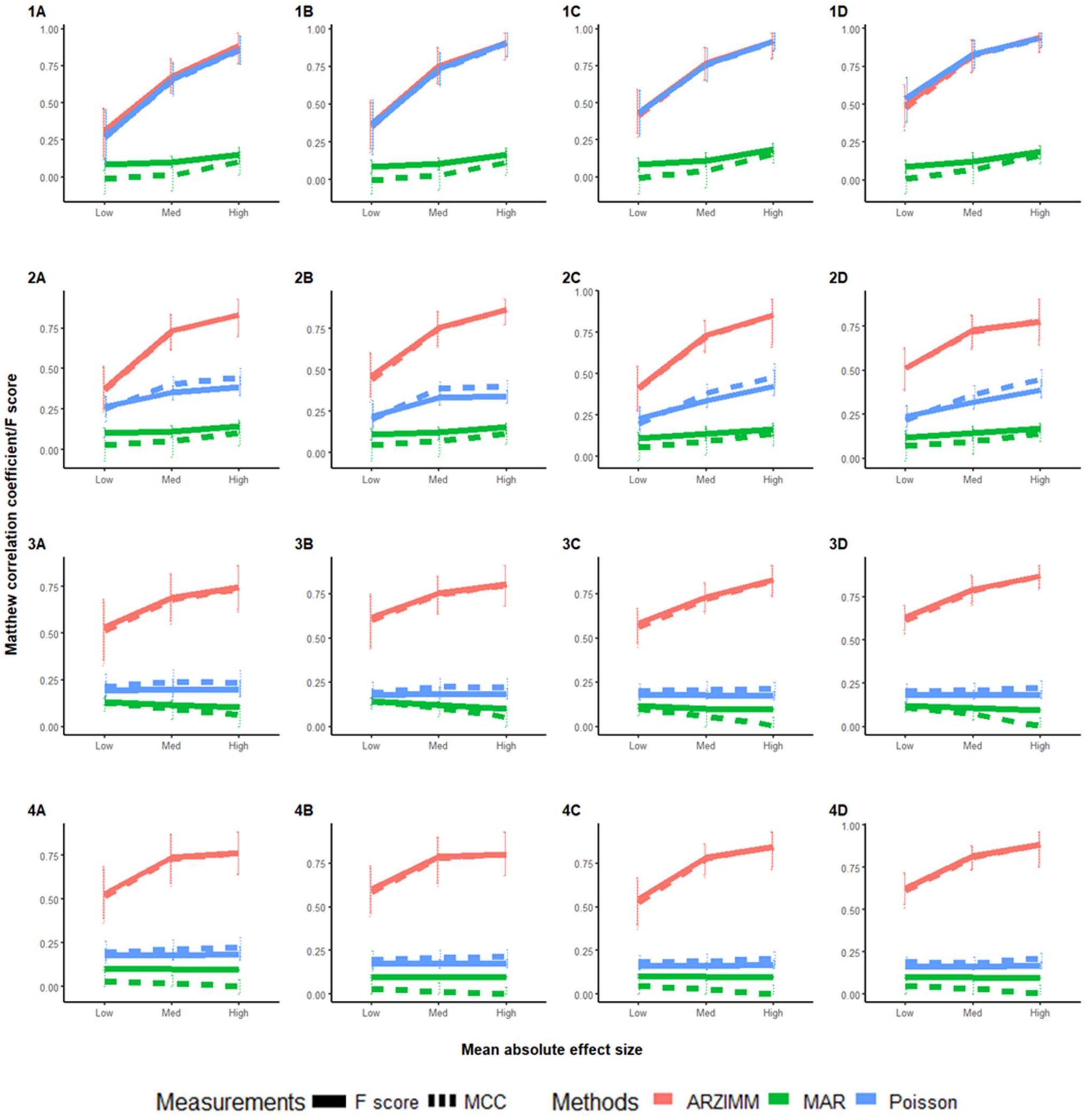


**Supplementary Figure 2.** Matthew correlation coefficient and F-score over 500 simulations for four scenarios for (A) 20 subjects with 10 time points; (B) 20 subjects with 10 time points; (C) 50 subjects with 10 time points; (D) 50 subjects with 20 time points. Poisson refers to the penalized Poisson autoregression model and MAR refers to penalized log-normal multivariate autoregression model. False Positive Rate and True Positive Rate are reported with mean and 95% interval over 500 simulations for four scenarios to have: (1) no zero-inflated structure and no heterogeneity; (2) heterogeneity but no zero-inflated structure; (3) zero-inflated structure but no heterogeneity; (4) both zero-inflated structure and heterogeneity.

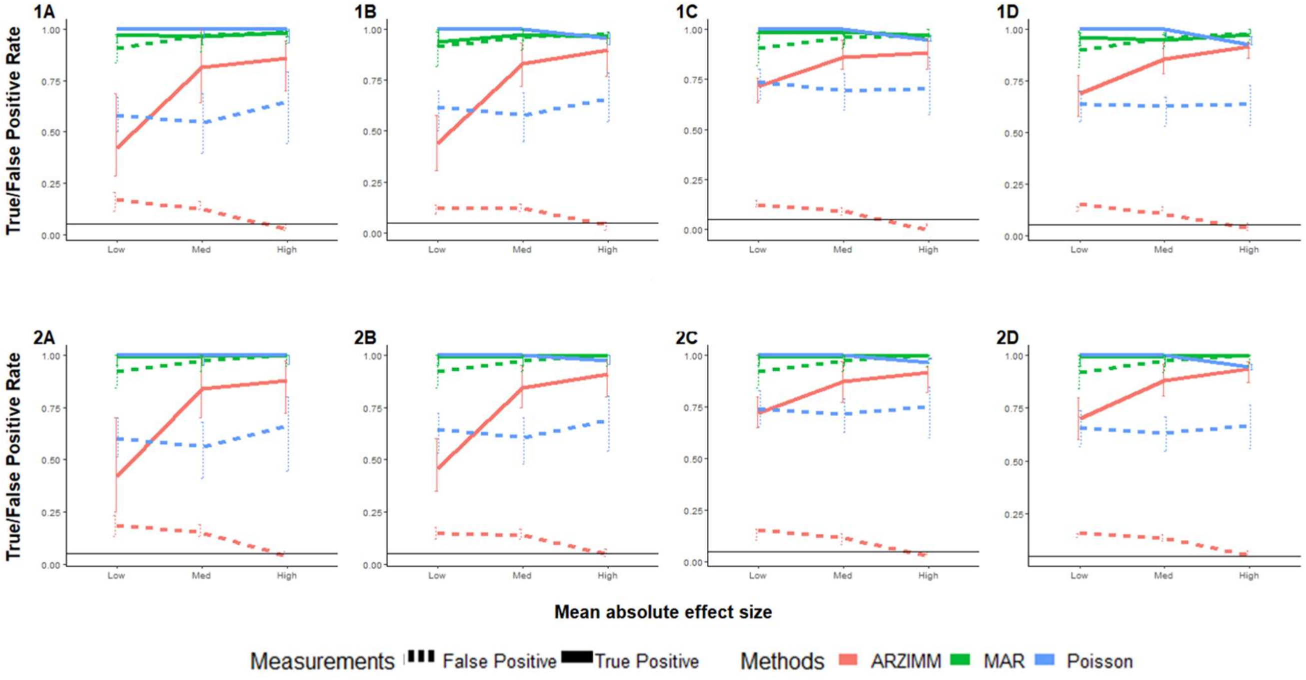


**Supplementary Figure 3.** Simulation results of mis-specified model for (A) 20 subjects with 10 time points; (B) 20 subjects with 20 time points; (C) 50 subjects with 10 time points; and (D) 50 subjects with 20 time points. Instead of true zero inflation proportions $\boldsymbol{p}_i$, the mis-specified zero inflation proportions $\widetilde{\boldsymbol{p}}_i = \boldsymbol{p}_i + \alpha\beta_i$ are used to simulate absolute abundances. Poisson refers to the penalized Poisson autoregression model and MAR refers to penalized log-normal multivariate autoregression model. False Positive Rate and True Positive Rate are reported with mean and 95% interval over 500 simulations for 3 scenarios to have $\alpha \sim Bernoulli(0.1)$ and $\beta_i$'s $m$-th element: (1) $\beta_{im} \sim N(0, 0.2p_{im})$ and (2) $\beta_{im} \sim N(0, 0.3p_{im})$.
